# Research on Intelligent Optimization of Farm Planting Strategies Driven by Crop Simulation Models: A Case Study of Farm X

**DOI:** 10.64898/2026.04.27.720996

**Authors:** Xinhang Lyu, Renjing Yu, Rongsheng Zhu

## Abstract

To meet the growing demand for precision and intelligent agricultural management, crop simulation models offer substantial potential for optimizing farm planting strategies. By simulating crop growth processes and assessing the effects of different management practices, these models provide a scientific basis for planting decision-making. In this study, the DSSAT model was first used to optimize the planting strategies of Farm X in 2023. Based on the optimized plans, the model was further applied to predict crop yields per unit area for 2024 and to establish the relationships among yield, planting density, and fertilizer application rate. Subsequently, SPSS was employed to develop a regression model describing the relationship among net profit per unit area, planting density, and fertilizer application rate. A genetic algorithm was then used to identify the optimal solutions under different scenarios, generating prescription maps for the optimal planting density and fertilizer application rate for each plot of Farm X in 2024. The results provide a scientific reference for the mechanized and automated implementation of field management practices and support the dual optimization of economic returns and resource use efficiency. This study not only conducted a systematic optimization of Farm X planting strategies for 2023, but also provided detailed predictions and optimized prescriptions for 2024 in a visual and practical form. The proposed approach offers a scientific decision-support tool for farm planting strategy formulation and lays a foundation for the intelligent and automated development of modern agriculture.

## 1 Introduction

Heilongjiang Province is the largest commercial grain production base in China, with both its grain commodity volume and commodity rate ranking first in the country [1]. Endowed with abundant water resources and extensive arable land, the province enjoys highly favorable natural conditions for agricultural production. Nevertheless, suboptimal planting density and fertilization practices have long constrained the yield potential of food crops in Heilongjiang Province. Therefore, determining appropriate planting density and fertilizer application rates to enhance crop productivity has become an issue of considerable urgency.

The DSSAT (Decision Support System for Agrotechnology Transfer) is a widely used crop simulation model system [2,3]. It can simulate crop growth and yield under different planting strategies [4] and evaluate the effects of various agricultural management practices on crop productivity and economic returns [5].

S et al. [6] reported that, in winter wheat production on the North China Plain, the DSSAT-CERES model was effective for evaluating nitrogen fertilizer management, improving nitrogen use efficiency, and reducing environmental pollution. Wang et al. [7] conducted a two-year experiment (2016–2017) in the High-Tech Agricultural Demonstration Park of Wugong County, Shanxi Province, and found that water and fertilizer stress significantly affected model sensitivity, while soil parameter calibration played an important role in improving simulation accuracy. Rugira et al. [8] applied the DSSAT maize model in the Loess Plateau region to determine suitable irrigation management practices and optimal sowing periods, thereby helping to stabilize spring maize production and improve regional water use efficiency. Zhang et al. [9] used data from the North China Plain together with the DSSAT-CERES-Wheat model to investigate the responses of winter wheat productivity and nitrogen use to planting density and nitrogen application rate under limited irrigation, providing a reference for the optimization of management strategies. Chen et al. [10] combined the DSSAT-CERES-Maize model with a dynamic irrigation algorithm in experiments conducted in northwestern China and found that maize yield predictions became progressively more stable before harvest. Qu et al. [11] used the DSSAT model to simulate changes in winter wheat yield in the Huang-Huai Plain under different climate scenarios, showing that increases in solar radiation and precipitation were beneficial to yield, whereas rising temperature had negative effects. Wei et al. [12] applied the DSSAT-CROPGRO-Soybean model to assess soybean drought risk under different irrigation levels in Bengbu, Anhui Province, and established a relationship curve between drought intensity and yield loss.

Internationally, research on and applications of the DSSAT crop simulation model have become relatively mature. As early as 1998, Jones et al. [13] proposed the integration of DSSAT into spatial decision support systems and outlined its framework, design principles, and major limitations. Anar et al. [14] modified the CERES-Sugarbeet model within DSSAT and incorporated it into the Cropping System Model (CSM) to simulate sugar beet growth, development, and yield. Mehrabi et al. [15] demonstrated that DSSAT could effectively predict wheat growth and yield under different semi-arid climatic conditions, irrigation strategies, planting methods, and nitrogen application rates. Their results showed that water stress and unfavorable climatic conditions significantly reduced grain yield, while exerting comparatively limited effects on straw yield. Jing et al. [16] applied the DSSAT wheat model to evaluate the effects of water management on spring wheat production in the Canadian Prairies and found that irrigation and drought-tolerant cultivars could improve both yield and water-use efficiency. Asgari et al. [17] calibrated and validated the DSSAT-CROPGRO-Rapeseed model using field experimental data and predicted the effects of climate change on the growth, yield, and water-use efficiency of winter rapeseed. Their study indicated that appropriate drainage management and agronomic practices could help achieve optimal yields under changing climatic conditions. Malik et al. [18] calibrated the DSSAT model to assess the impacts of agricultural management practices on irrigation requirements and nitrogen losses in intensively irrigated regions of Spain, thereby supporting the optimization of both grain yield and environmental performance. Similarly, Mehrabi et al. [19] used DSSAT to predict wheat yields in semi-arid regions and confirmed that water stress and climatic unsuitability were major factors limiting yield, while having relatively little influence on straw production. Hasan et al. [20] employed DSSAT to analyze the effects of climate change on rice yield in Bangladesh and identified temperature and rainfall patterns as key determinants; their results further showed that adjusting transplanting dates could mitigate some of the adverse effects of climate change. Pokhrel et al. [21] simulated energy cane growth using the DSSAT-CANEGRO model and found that increased irrigation significantly enhanced biomass production, providing a basis for evaluating the cultivation potential of sugarcane in semi-arid regions. Balpande et al. [22] investigated the effects of planting date on potato yield and reported that timely planting reduced yield variability, whereas delayed planting increased yield uncertainty.

In summary, previous studies have confirmed the strong capability of DSSAT in crop growth simulation, yield prediction, and management strategy evaluation. However, most existing research has concentrated on irrigation, nitrogen application, planting date, or climate factors separately, and less attention has been paid to the coupled optimization of planting density and fertilizer application under spatially heterogeneous field conditions. Moreover, although DSSAT has been widely used to assess agronomic and environmental outcomes, its integration with economic analysis and prescription-based decision-making for precision agriculture remains limited. Therefore, it is necessary to further explore the combined use of DSSAT and optimization approaches to determine optimal management strategies and provide a scientific basis for high-yield, high-efficiency, and precise agricultural production.

This study takes Farm X in Heilongjiang Province as the research object and seeks to develop a scientifically grounded and practical planting strategy through detailed analysis and prediction. Centering on DSSAT-based simulation and optimization of intelligent planting strategies for X Farm [23], the main contributions of this paper are as follows:

1. A system-level analysis of X Farm was carried out, revealing substantial room for optimization in its current production management.
2. The DSSAT model was employed to investigate the effects of yield-related factors during the crop growth period on final yield at X Farm.
3. Based on the DSSAT model, the planting strategy of X Farm was further optimized to improve management efficiency and support precise agricultural decision-making.

The remainder of this paper is organized as follows. Section 2 describes the experimental methods. Section 3 presents the experimental results. Section 4 discusses the findings and the limitations of the proposed approach. Finally, Section 5 concludes the paper.

## 2 Materials and Methods

### 2.1 Experimental Location

Farm X is situated in Suibin County, Hegang City, Heilongjiang Province. Characterized by fertile black soil and a temperate monsoon climate, the region provides highly favorable natural conditions for agricultural production. The farm comprises 54 plots with distinct characteristics, numbered from 1 to 54. Among these, plots 10–21 and 36–47 are allocated to paddy rice cultivation, whereas the remaining plots are classified as dryland fields, primarily used for maize and soybean production. The topographic map of the study area is shown in Figure 1. Figure 2 presents the spatial layout of the plot numbers within Farm X, and Figure 3 shows the area of each plot in hectares.

**Figure 1.**
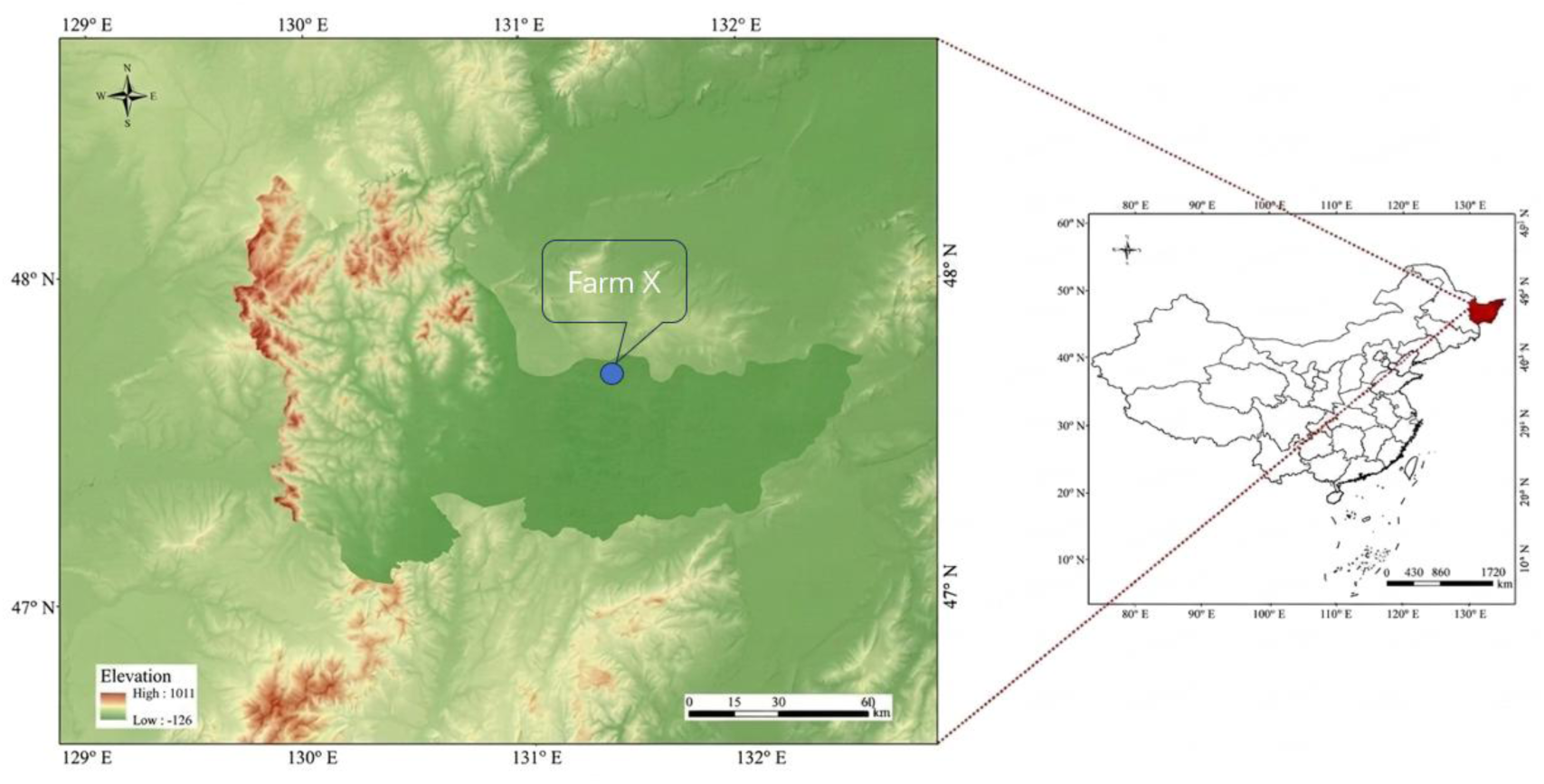
Topographic map of the area where the Farm X is located.

**Figure 2.**
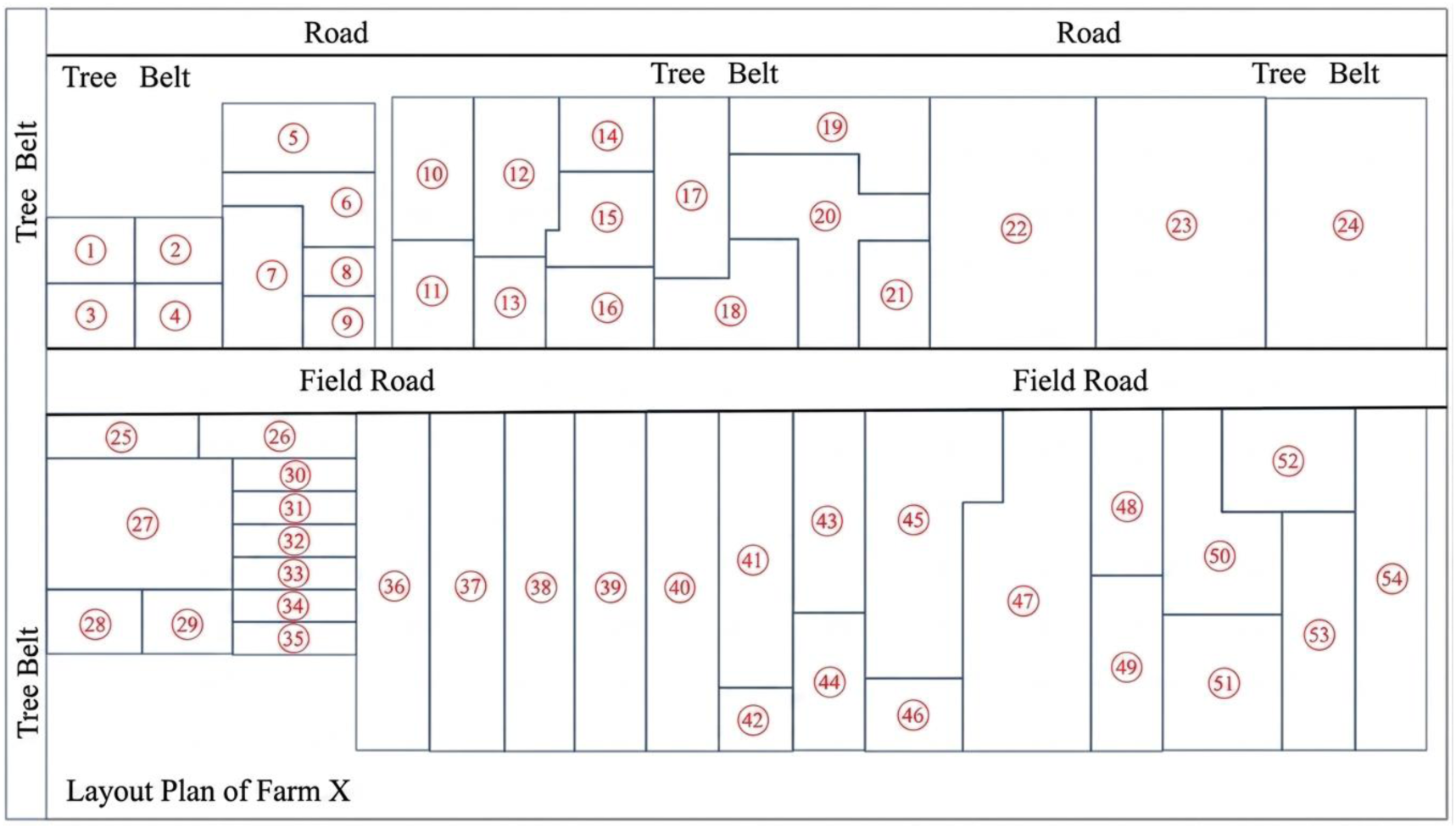
Plan diagram of numbering for each plot of the Farm X.

**Figure 3.**
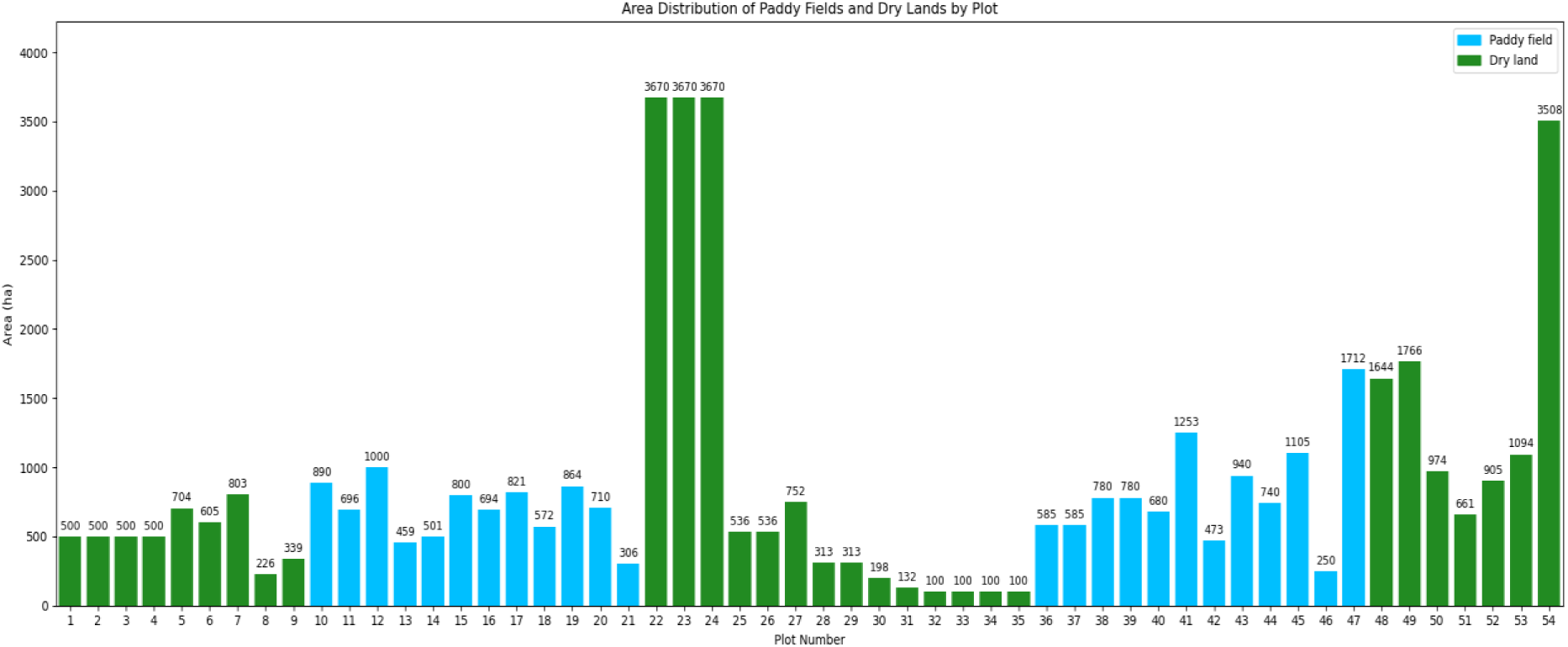
Area map of each plot of the Farm X (ha)

### 2.2 Technical Roadmap

The provided flowchart illustrates a comprehensive research methodology for optimizing agricultural planting strategies through a combination of crop growth simulation, economic forecasting, and heuristic optimization. The research framework is structured as follows:

#### 1) Model Calibration and Data Integration

The process begins with the collection of multi-source environmental and agronomic data, including meteorological data, varietal genetic parameters, field management data, and soil data. These inputs are utilized to characterize the "Original planting strategy of X Farm." This data forms the basis for the localization and calibration of the Decision Support System for Agrotechnology Transfer (DSSAT) model, ensuring that the biophysical simulations are tailored to the specific conditions of the study site.

#### 2) Simulation of Planting Schemes

In parallel, the study identifies the three major cultivated crops relevant to the farm. By varying management inputs and environmental conditions, the researchers design 36 interactive planting schemes. These schemes are processed through the calibrated DSSAT model to generate a robust dataset of predicted yields.

#### 3) Regression and Economic Modeling

To bridge the gap between biophysical simulation and economic optimization, the workflow proceeds in two directions:

Yield Modeling: A regression model is established based on the DSSAT output to provide a computationally efficient functional relationship between input variables and predicted yields.

Economic Analysis: Using historical economic data of agricultural products, the study performs a prediction of selling prices and costs.

#### 4) Optimization Framework

These two streams converge to construct a net profit optimization model specifically for Farm X. The objective function seeks to maximize profitability while considering the constraints defined by the regression model and economic forecasts.

Given the complexity of the solution space, the study employs a Genetic Algorithm (GA) to solve the optimization problem. The GA iteratively evolves potential solutions to identify the most effective configurations.

### 2.3 DSSAT Model introduction and localization

#### 2.3.1 Introduction to the DSSAT Model

The DSSAT model (Decision Support System for Agrotechnology Transfer) is a comprehensive software suite designed to support crop management decision-making and improve agricultural production efficiency [24]. It integrates a series of process-based simulation modules that accurately represent crop growth and development [25], allowing the simulation of crop production under diverse climatic conditions, soil types, and crop varieties. The DSSAT software package can simulate 16 different crops, including rice, maize, and soybean, with this study focusing specifically on these three crops. To ensure that the simulated outputs closely match observed field data, local calibration of model parameters using experimental datasets is essential during model application [26].

#### 2.3.2 Localization of DSSAT Model

Prior to applying the DSSAT model to Farm X, it is necessary to calibrate and validate the genetic parameters of the crop varieties grown on the farm to ensure consistency with local conditions. To improve the accuracy and reliability of the model outputs [27], these varietal parameters should be systematically adjusted, tested, and verified. In DSSAT, this iterative calibration and validation process can be carried out using built-in tools such as the GLUE (Generalized Likelihood Uncertainty Estimation) function [28].

In this study, the genetic parameters of maize (Demeiya 3), rice (Longjing 1624), and soybean (Kendou 94) varieties were obtained using the above-mentioned methods, and the consistency [29] between the model predictions and the observed data (emergence period, flowering period, maturity period, and yield per unit area) was evaluated. The parameter adjustment verification results of the simulated and measured values of maize yield in each plot of Farm X in 2023 based on the DSSAT model are shown in Figure 5. The parameter adjustment verification results of the simulated and measured values of rice yield in each plot of Farm X in 2023 based on the DSSAT model are shown in Figure 6. The parameter adjustment verification results of the simulated and measured soybean yield of each plot in Farm X in 2023 based on the DSSAT model are shown in Figure 7.

**Figure 4.**
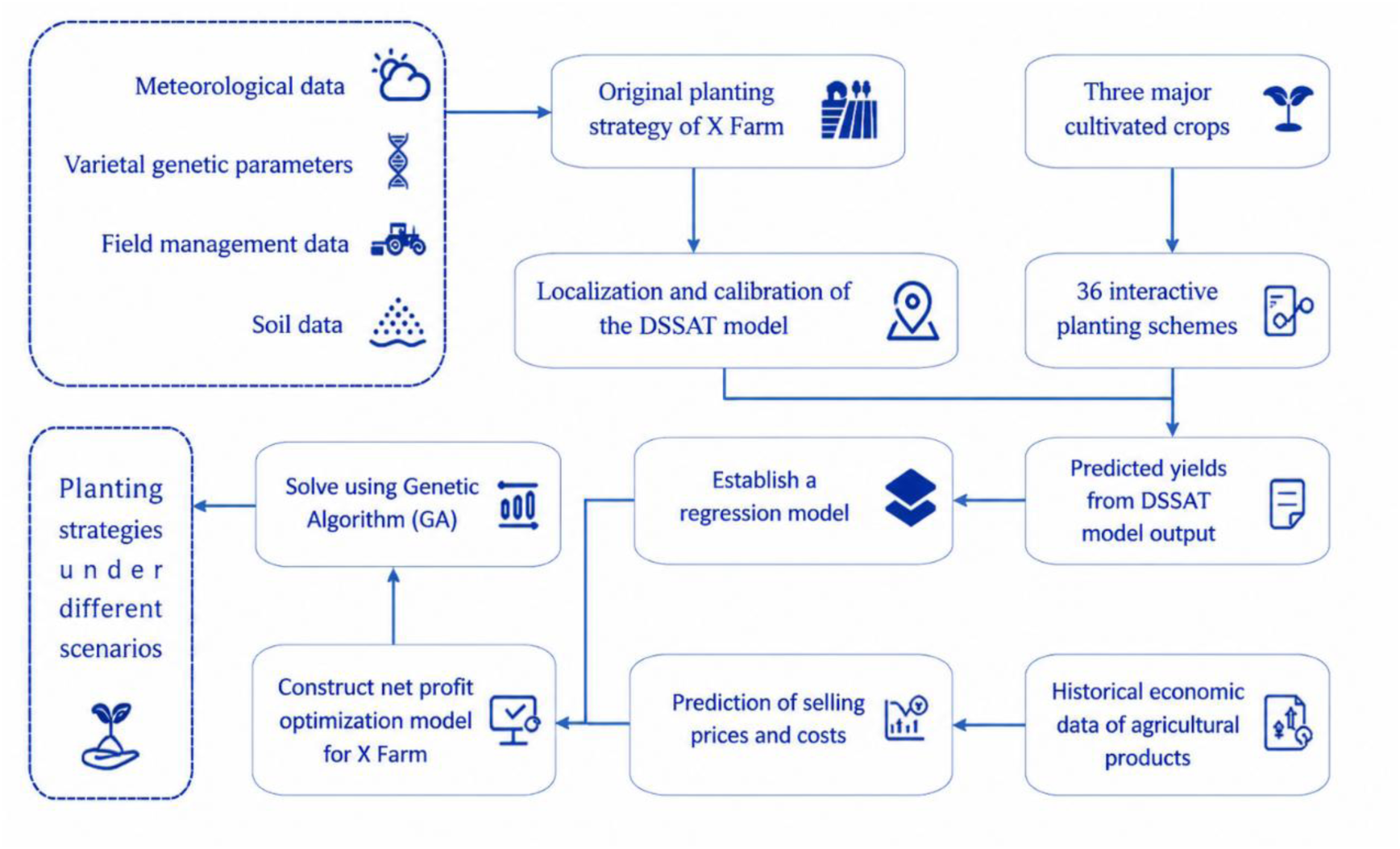
Technology route.

**Figure 5.**
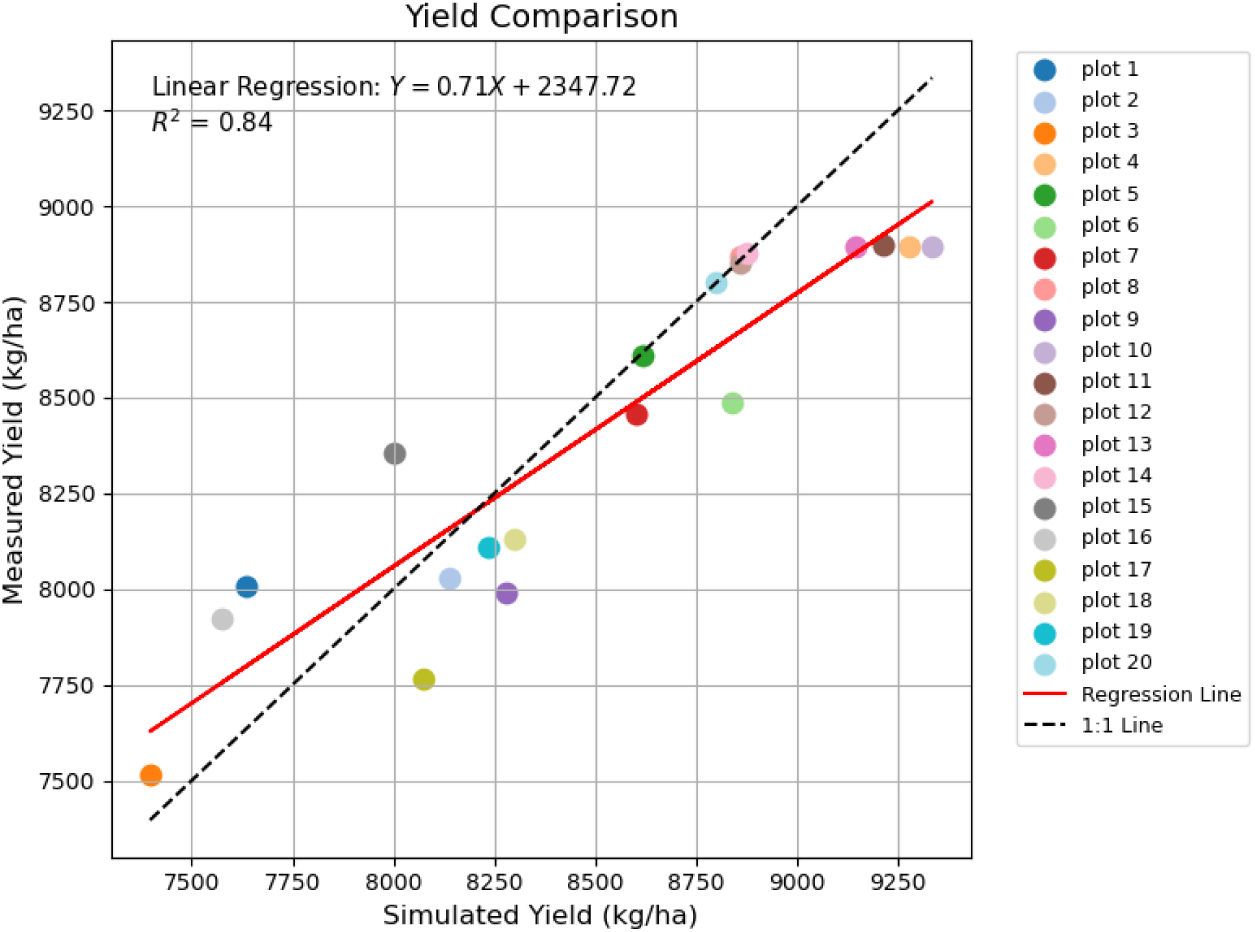
Verification results of simulated and measured maize yield values for each plot of the Farm X in 2023 based on DSSAT model.

**Figure 6.**
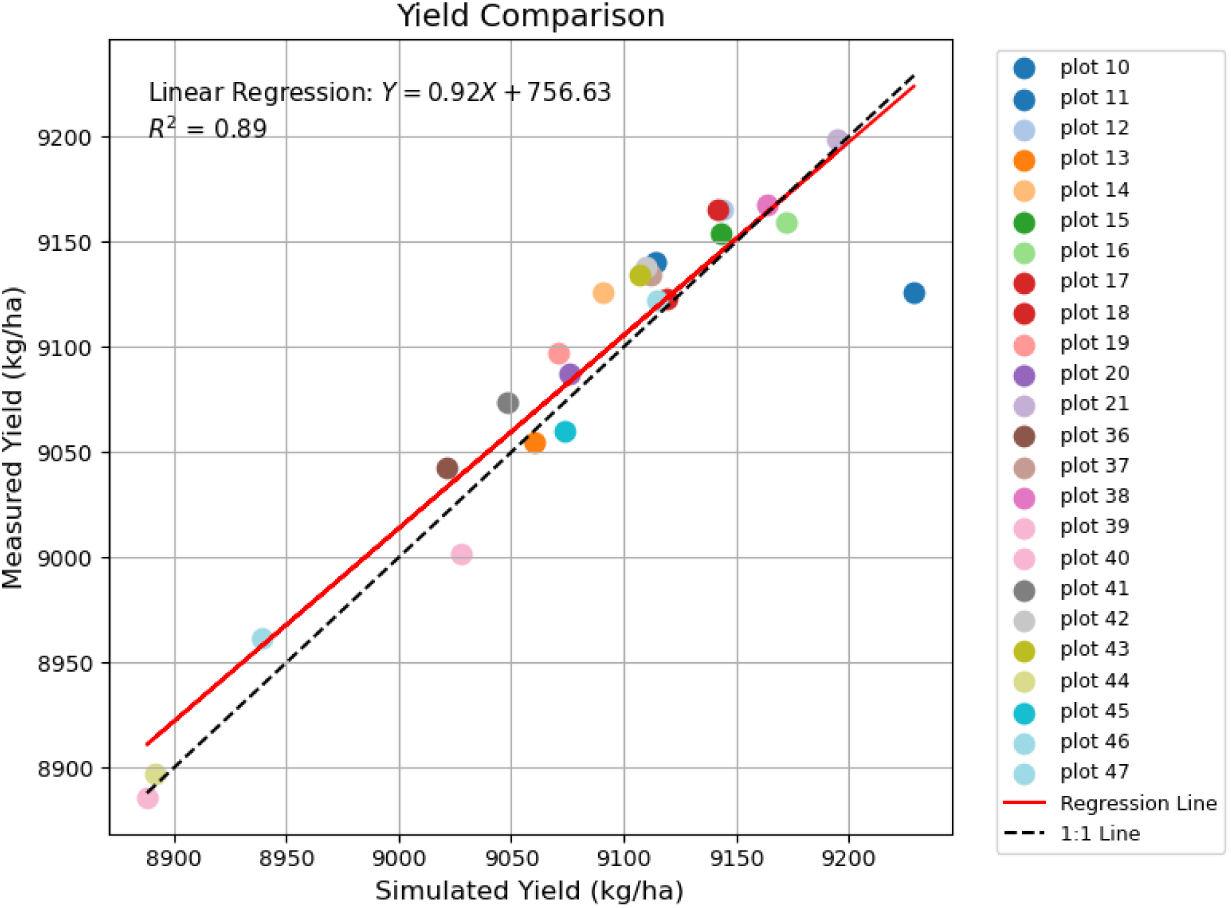
Verification results of simulated and measured rice yield values for each plot of the Farm X in 2023 based on DSSAT model.

**Figure 7.**
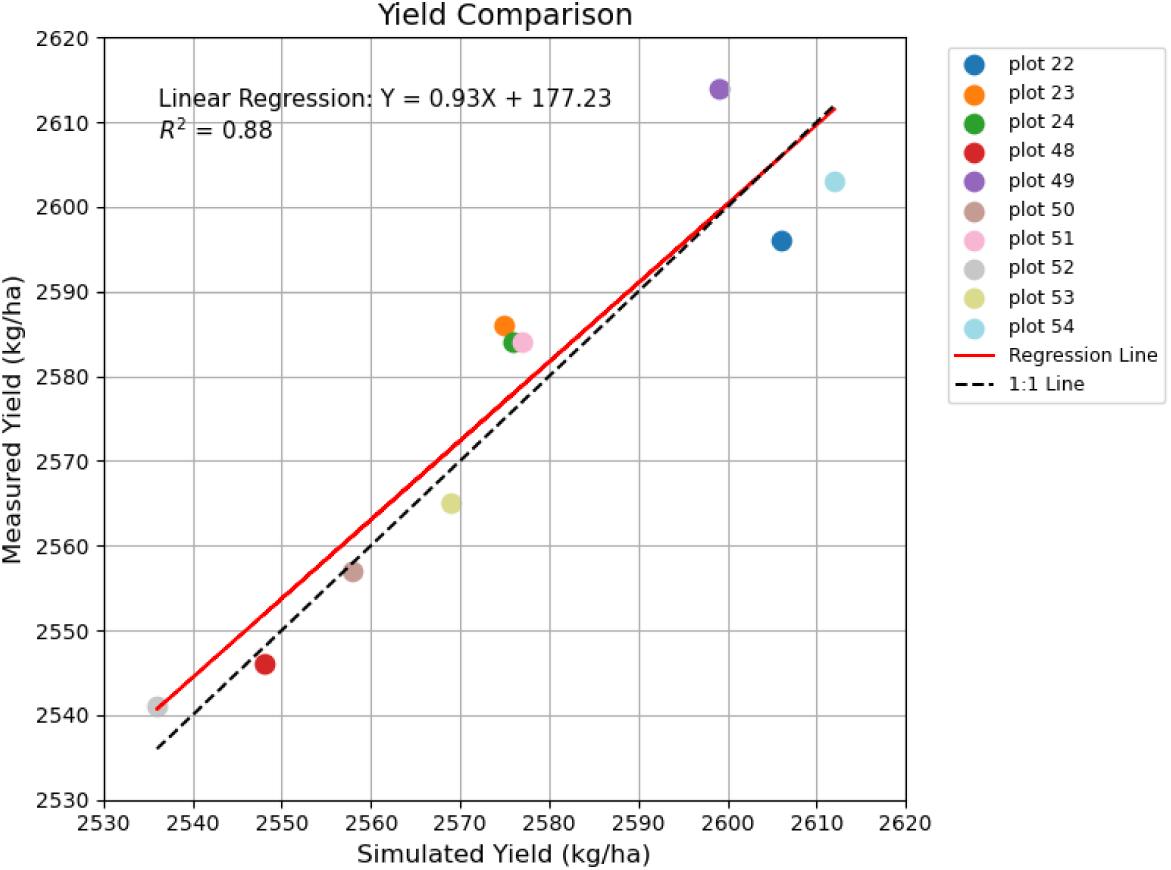
Verification results of simulated and measured soybean yield values for each plot of the Farm X in 2023 based on DSSAT model.

### 2.4 Analysis of X Farm Planting Strategies Based on DSSAT Model

#### 2.4.1 Optimized Planting strategy

This study, based on the DSSAT model, set up 36 different intercropping schemes with six different fertilization treatments and six different planting densities as mentioned above to simulate and predict the crop ^[30]^ (rice, maize and soybean) yields of various plots in Farm X, Heilongjiang Province in 2024. The main crop varieties grown in various plots of Farm X are still maize (Demeiya 3), rice (Longjing 1624), and soybeans (Kendou 94). Sowing, irrigation and soil mounding, etc. are set up according to the actual situation of Farm X, and compound fertilizer (N: P: K=15% : 15% : 15%) is used as fertilizer.

The planting density of rice (Longjing 1624) is set at six types, namely:15 plant/m^2^ (P1), 20 plant/m^2^ (P2), 25 plant/m^2^ (P3), 30 plant/m^2^ (P4), 35 plant/m^2^ (P5),40 plant/m^2^ (P6).

The fertilizer application rates for rice (Longjing 1624) are set at six types, namely:280kg/ha(S1),320kg/ha(S2),360kg/ha(S3),400kg/ha(S4),440kg/ha(S5),480 kg/ha (S6). The planting density of maize (Demeiya No. 3) is set at six types, namely: 3 plant/m^2^ (P1), 6 plant/m^2^ (P2), 9 plant/m^2^ (P3), 12 plant/m^2^ (P4), 15 plant/m^2^ (P5), 18 plant/m^2^ (P6). The fertilization rate for maize (Demeiya No. 3) is set at six types, namely:140 kg/ha (S1),180 kg/ha (S2),220 kg/ha (S3),260 kg/ha (S4),300 kg/ha (S5),340 kg/ha (S6). The planting density of soybeans (Kendou 94) is set at six types, namely:10 plant/m^2^ (P1),15 plant/m^2^ (P2),20 plant/m^2^ (P3),25 plant/m^2^ (P4),30 plant/m^2^ (P5),35 plant/m^2^ (P6). The fertilizer application rates for soybeans (Kendou 94) are set at six types, namely:40 kg/ha (S1),80 kg/ha (S2),120 kg/ha (S3),160 kg/ha (S4),200 kg/ha (S5),240 kg/ha (S6).

Farm X has set up a total of 36 interactive planting schemes based on the planting density and fertilizer application amount of each crop mentioned above. The planting plan of Farm X includes a total of 36 different interactive planting plans, as shown in

**Table 1.** 36 different interactive planting schemes for the Farm X.

| Plan | Density<br>P1 | Density<br>P2 | Density<br>P3 | Density<br>P4 | Density<br>P5 | Density<br>P6 |
| --- | --- | --- | --- | --- | --- | --- |
| Fertilizer<br>S1 | P1S1 | P2S1 | P3S1 | P4S1 | P5S1 | P6S1 |
| Fertilizer<br>S2 | P1S2 | P2S2 | P3S2 | P4S2 | P5S2 | P6S2 |
| Fertilizer<br>S3 | P1S3 | P2S3 | P3S3 | P4S3 | P5S3 | P6S3 |
| Fertilizer<br>S4 | P1S4 | P2S4 | P3S4 | P4S4 | P5S4 | P6S4 |
| Fertilizer<br>S5 | P1S5 | P2S5 | P3S5 | P4S5 | P5S5 | P6S5 |
| Fertilizer<br>S6 | P1S6 | P2S6 | P3S6 | P4S6 | P5S6 | P6S6 |

#### 2.4.2 Analysis of the unit net profit of Crops Grown in Each plot of Farm X Based on the DSSAT model

This study established experiments based on 36 distinct interactive planting schemes, which varied in planting density and fertilization amounts, without altering the crop varieties cultivated in the plots of X Farm in 2023. Employing the DSSAT model, a comprehensive analysis and simulation of unit crop yields for each plot was conducted [31]. Furthermore, this research considered various factors, including planting costs, to ascertain the unit net profit of crops across the plots of X Farm under the 36 different interactive planting schemes. This investigation not only examined the interactive planting effects among different crops but also provided a thorough analysis of the planting schemes to optimize the benefits and efficiency of the farmland [32].]。 The highest unit net income of crops planted in each plot of Farm X in 2023 simulated by the DSSAT model is shown in Table 2.

**Table 2.** The highest unit net income of crops planted on each plot of the Farm X in 2023 simulated by DSSAT model (yuan/ha)

| **N1** | **N2** | **N3** | **N4** | **N5** | **N6** | **N7** | **N8** | **N9** |
| --- | --- | --- | --- | --- | --- | --- | --- | --- |
| Maize<br>13892 | Maize<br>11975 | Maize<br>12167 | Maize<br>14583 | Maize<br>13658 | Maize<br>12089 | Maize<br>12520 | Maize<br>14044 | Maize<br>11967 |
| **N10** | **N11** | **N12** | **N13** | **N14** | **N15** | **N16** | **N17** | **N18** |
| Rice<br>11008 | Rice<br>11017 | Rice<br>11044 | Rice<br>11000 | Rice<br>11019 | Rice<br>11319 | Rice<br>10708 | Rice<br>11031 | Rice<br>11028 |
| **N19** | **N20** | **N21** | **N22** | **N23** | **N24** | **N25** | **N26** | **N27** |
| Rice<br>11019 | Rice<br>11017 | Rice<br>11061 | Soybean<br>9152 | Soybean<br>9221 | Soybean<br>9129 | Maize<br>14640 | Maize<br>14370 | Maize<br>14533 |
| **N28** | **N29** | **N30** | **N31** | **N32** | **N33** | **N34** | **N35** | **N36** |
| Maize<br>14211 | Maize<br>14783 | Maize<br>12311 | Maize<br>11936 | Maize<br>11326 | Maize<br>12692 | Maize<br>12407 | Maize<br>14401 | Rice<br>10648 |
| **N37** | **N38** | **N39** | **N40** | **N41** | **N42** | **N43** | **N44** | **N45** |
| Rice<br>11039 | Rice<br>11036 | Rice<br>10790 | Rice<br>10405 | Rice<br>10965 | Rice<br>11014 | Rice<br>11036 | Rice<br>10468 | Rice<br>11011 |
| **N46** | **N47** | **N48** | **N49** | **N50** | **N51** | **N52** | **N53** | **N54** |
| Rice<br>10129 | Rice<br>10993 | Soy<br>9216 | Soy<br>9239 | Soy<br>9204 | Soy<br>9359 | Soy<br>9164 | Soy<br>9170 | Soy<br>9198 |

The simulation results indicate that in 2023, the net income from planting maize at farm X will be the lowest at 11,326 (yuan/ha), while the net income from rice will be 10,129 (yuan/ha), and from soybeans, it will be 9,129 (yuan/ha). Conversely, the highest net income from maize cultivation is projected to be 14,783 (yuan/ha), from rice 11,319 (yuan/ha), and from soybeans 9,359 (yuan/ha). A comparative analysis reveals that optimizing planting density and fertilization can enhance both crop yield and unit net income. Consequently, the farm should adapt its strategies to reflect the actual conditions. The existing planting strategy presents significant opportunities for improvement, thereby establishing a foundation for future research. By integrating the predictive outcomes of the DSSAT model, the formulation of more effective planting strategies could lead to increased farm profitability.

#### 2.4.3 Construct a regression model between the crop unit yield, planting density and fertilization amount in each plot of X farm

This chapter employs the DSSAT model to simulate and predict crop unit yields for each plot of farm X in 2024, utilizing 36 distinct combinations of planting density and fertilizer application. Consequently, crop unit yield data were obtained for these 36 combinations. To further investigate the relationship among crop unit yield, planting density, and fertilization amount, this study employs SPSS to establish a regression model. In SPSS, the unit yield of the planted crops in each plot of Farm X can be regarded as the dependent variable, while the planting density and fertilizer application amount can be taken as independent variables. A regression model can be constructed to analyze the relationship between them [30]. By analyzing the coefficients of the regression model, significance tests, the fit degree of the model and other statistical information, the extent and direction of the influence of planting density and fertilizer application on crop yield can be understood. By analyzing the results of the regression model, the contribution of planting density and fertilizer application to crop yield can be evaluated, and the planting strategy for optimizing yield can be identified. This will provide important references for farm managers to make reasonable planting decisions, optimize crop yields and resource utilization.

#### 2.4.4 Regression Model of crop unit yield per plot in Farm X Based on DSSAT Model and Its solution

This chapter’s research focuses on various plots of Farm X in 2024. Based on the 36 different planting densities and fertilizer application combinations mentioned earlier, an experimental plan was set up, and the DSSAT model was used to simulate and predict the unit yield of each plot. In Farm X, Plots 1 to 9, 22 to 35 and 48 to 54 are dry fields. On these dry fields, two types of dry field crops can be chosen to be grown, namely maize or soybeans. Plots No. 10 to No. 21 and No. 36 to No. 47 are paddy fields, and only rice can be chosen as the planting crop for these paddy fields. Based on obtaining the simulation prediction results of the unit yield of crops planted in each plot of Farm X in 2024, the function of establishing regression models in SPSS 25.0 software was used for analysis and a binary quadratic regression model was established with the unit yield (Y) as the predicted dependent variable and the fertilizer application amount (X_1_) and planting density (X_2_) as independent variables. By developing a regression model that relates crop unit yield to fertilization amount and planting density for each plot of X farm, we can predict the maximum unit yield of various crops across different plots in 2024, along with the corresponding fertilization amounts and planting densities. This information will assist farm decision-makers in evaluating and optimizing their planting strategies.

To gain a deeper understanding of the patterns behind these data, this paper employed SPSS statistical analysis, taking the unit yield (Y) of crops in each plot of Farm X as the predicted dependent variable, and the amount of fertilizer application (X_1_) and planting density (X_2_) as the key independent variables. Through regression analysis, a statistical model was established between the unit yield of crops on Farm X in 2024 and the planting density and fertilizer application amount:

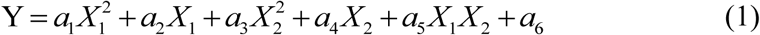

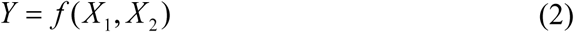

Perform derivative operations on formula (1) to optimize and obtain the optimal

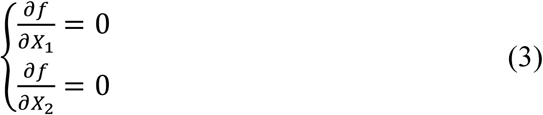

Obtain the optimal solution of fertilizer application amount (*X*_1_) and planting density (*X*_2_).

Taking the maize planting in Plot 1 of Farm X in 2024 as an example, a binary quadratic regression model for maize planting in Plot 1 was obtained through SPSS analysis, where is the unit yield of maize(kg/ha), *X*_1_ is the fertilizer application amount(kg/ha) and *X*_2_ is the planting density(plant/m^2^), The equation is shown in formula (4).

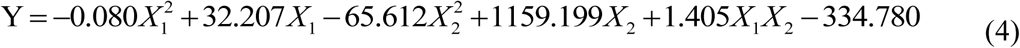

Through analysis, this study found that the P-value of this regression model is less than 0.01, while the coefficient of determination is greater than 0.900, and the F-value is 215.274, indicating that the model has extremely significant meaning and a good fit. Perform derivative operations on formula (4) to optimize and obtain the optimal solution of its extreme value:

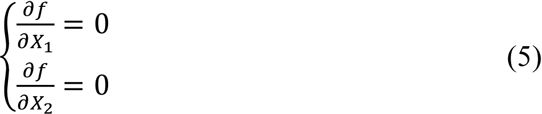

The optimal amount of fertilizer application is obtained by solving (*X*_1_) and planting density (*X*_2_):

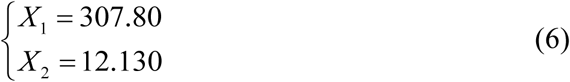

Substituting the calculated values of X1 and X2 into the regression equation (4) yields the Y value, representing the maximum unit yield of maize cultivated in Plot 1 of Farm X in 2024. With a fertilizer application of 307.80 kg/ha and a planting density of 12.130(plants/m^2^), the unit yield of maize attains its maximum at 11,652.13kg/ha. Concurrently, a three-dimensional map was generated using Origin 9.1 to illustrate the variations in yield for maize planted in Plot 1 of Farm X.

**Figure 7.**
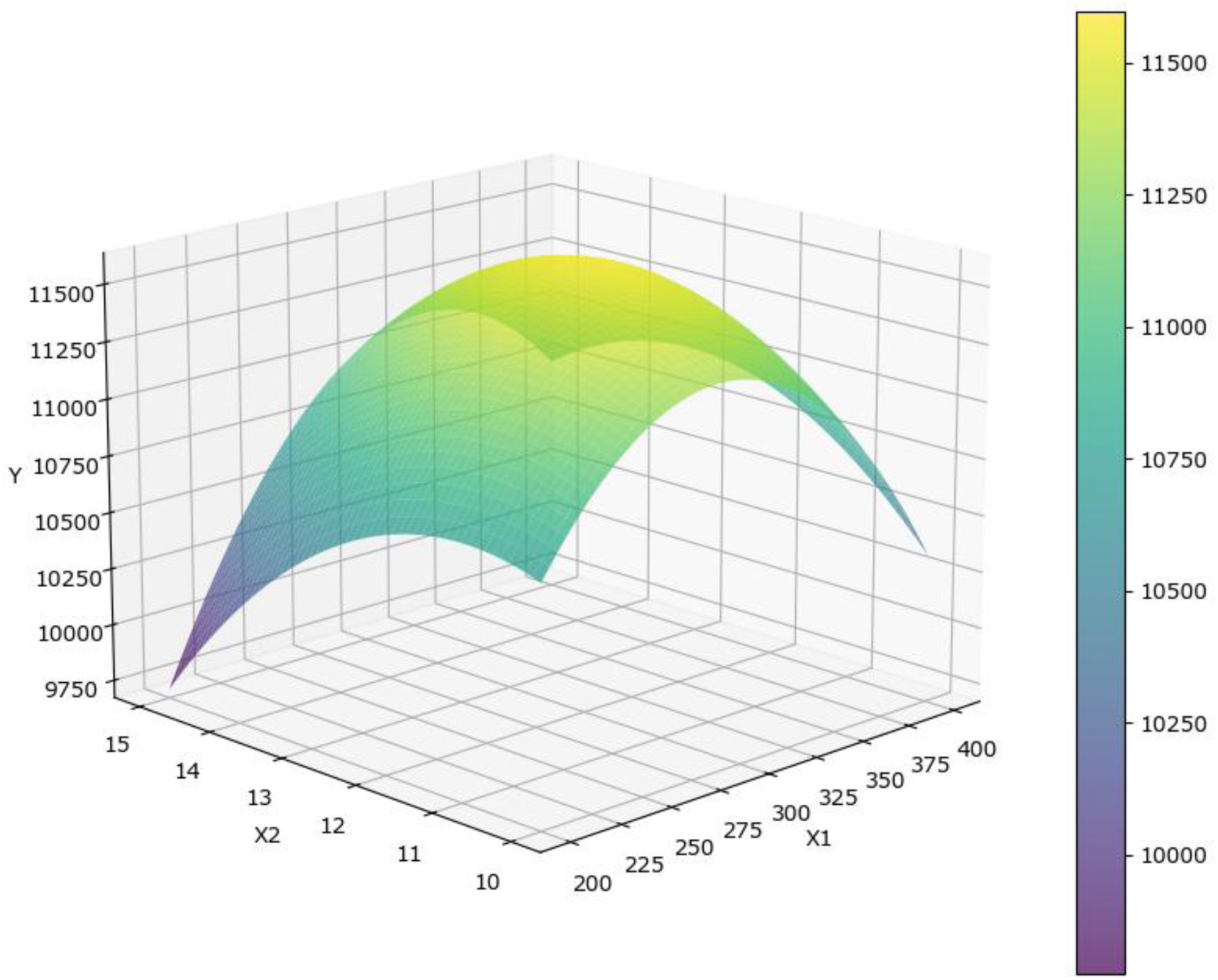
The unit yield of maize planted on plot 1 of the Farm X in 2024 varies with planting density and fertilization amount.

According to the above method, the regression models and optimal solutions for the unit yield of each crop were solved. The results are as follows: The regression models for the unit yield of maize crops planted in each plot of Farm X in 2024 are shown in **Appendix 1**. The optimal fertilizer application rate and planting density for maize crops planted in each plot and the corresponding unit yield are shown in **Appendix 2**. The regression model of the unit yield of rice crops planted in each plot of Farm X in 2024 is shown in **Appendix 3**. The optimal fertilizer application rate and planting density of rice crops planted in each plot and the corresponding unit yield are shown in **Appendix 4**. The regression model of the unit yield of soybean crops planted in each plot of Farm X in 2024 is shown in **Appendix 5**. The optimal fertilizer application rate, planting density and corresponding unit yield of soybean crops planted in each plot are shown in **Appendix 6**.

### 2.5 Research on Optimization of Planting Strategies for X Farm Based on DSSAT Model

#### 2.5.1 Use regression analysis methods to predict the time series of selling prices and planting costs of each crop

The historical sales prices and planting costs for each crop pertinent to the research in this chapter are sourced from the "Compilation of National Agricultural Products Cost and Benefit Data (2004-2023)." This compilation encompasses the sales price of each crop (yuan/kg), as well as the costs associated with fertilizers (yuan/ha), materials and services (yuan/ha), and labor (yuan/ha) necessary for the experiment. This study employs the time series forecasting function available in SPSS 25.0 software, integrating the sales price and planting cost data from previous years to project the selling price and planting cost of each crop for 2024. The modelling prediction results and associated statistics for maize planting costs at Farm X in 2024 are presented in Table 2. Similarly, the modelling prediction results and statistics for rice planting costs at Farm X in 2024 are detailed in Table 3. Finally, the modelling prediction results and statistics for soybean planting costs at Farm X in 2024 are illustrated in Table 4.

**Table 2.**
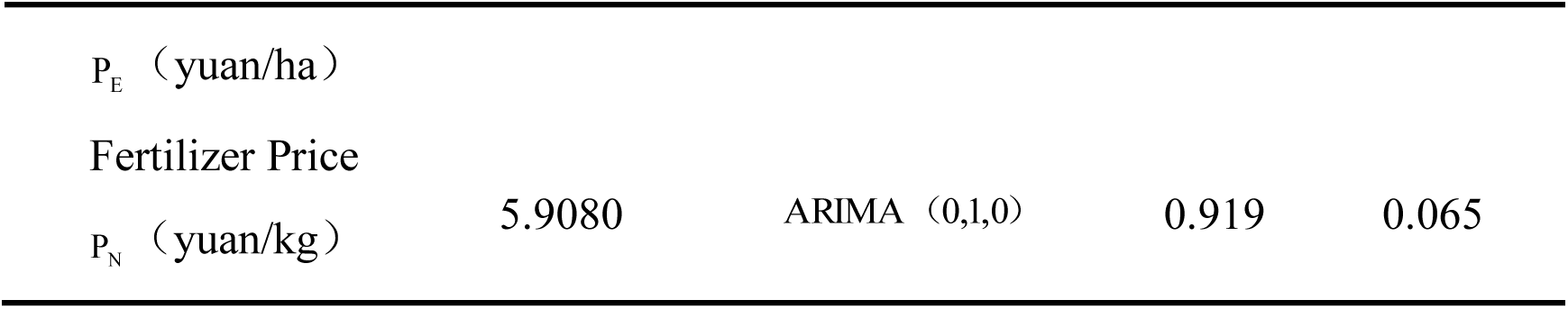
Modeling and prediction results and model statistics of maize planting costs in the Farm X in 2024.

**Table 3.** Modeling and prediction results and model statistics of rice planting costs in the Farm X in 2024.

| | Predictive Value | ARIMA ($p,d,q$) | $R^2$ | Significance |
| --- | --- | --- | --- | --- |
| Rice Price |  |  |  |  |
| $P_i$(yuan/kg) | 2.8366 | ARIMA (0,1,0) | 0.904 | 0.010 |
| Goods and services expenses |  |  |  |  |
| $P_{MS}$ (yuan/ha) | 6896.6906 | ARIMA (0,1,0) | 0.922 | 0.014 |
| Labor cost |  |  |  |  |
| $P_E$ (yuan/ha) | 7352.8474 | ARIMA (0,1,0) | 0.967 | 0.006 |
| Fertilizer Price |  |  |  |  |
| $P_N$ (yuan/kg) | 6.8960 | ARIMA (0,1,0) | 0.914 | 0.098 |

**Table 4.**
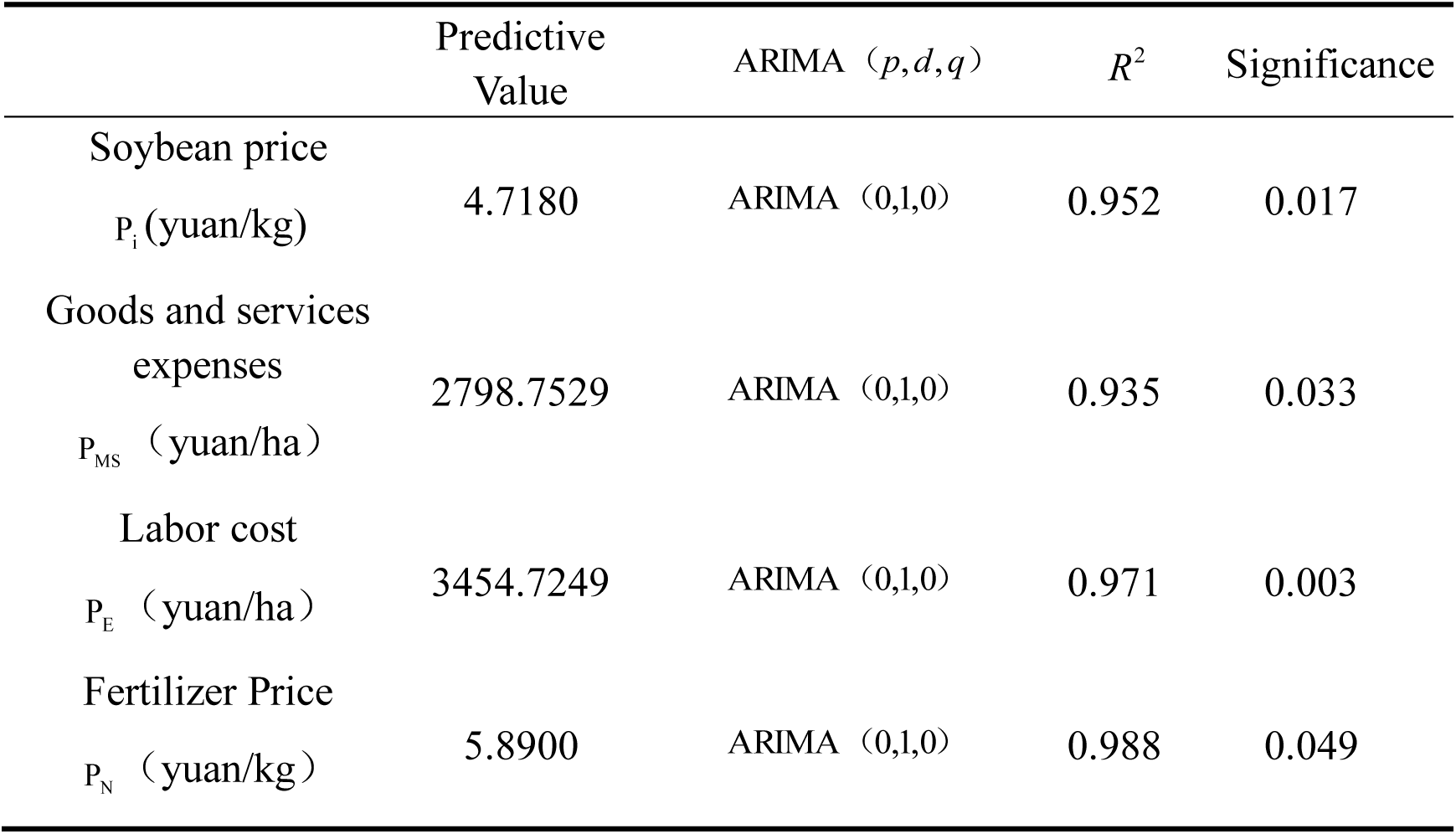
Modeling and prediction results and model statistics of soybean planting costs in the Farm X in 2024.

| | Predictive Value | ARIMA ($p,d,q$) | $R^2$ | Significance |
| --- | --- | --- | --- | --- |
| Soybean price |  |  |  |  |
| $P_i$(yuan/kg) | 4.7180 | ARIMA (0,1,0) | 0.952 | 0.017 |
| Goods and services expenses |  |  |  |  |
| $P_{MS}$ (yuan/ha) | 2798.7529 | ARIMA (0,1,0) | 0.935 | 0.033 |
| Labor cost |  |  |  |  |
| $P_E$ (yuan/ha) | 3454.7249 | ARIMA (0,1,0) | 0.971 | 0.003 |
| Fertilizer Price |  |  |  |  |
| $P_N$ (yuan/kg) | 5.8900 | ARIMA (0,1,0) | 0.988 | 0.049 |

This chapter employs cost-benefit analysis to determine the net profit of crops cultivated on the farm. Input costs encompass the expenses associated with planting each crop, which include fertilisation costs, material and service expenses (notably, the seed fee for each plot remains constant regardless of planting density), and labour costs. The net profit derived from crop cultivation in each plot is calculated as the gross income from each crop minus the total input costs.

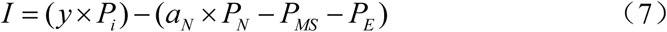

In Formula (2-4), *I* represents the net income of each plot in Farm X, *y* represents the yield of each crop in each plot, *P*_*i*_ represents the market selling price of the crops, *a*_*N*_represents the amount of fertilizer applied, *P*_*N*_ represents the price of fertilizers, *P*_*MS*_represents the cost of materials and services, and *P*_*E*_represents the labor cost. *y* × *P*_*i*_ represents the gross income of each crop in each plot, and *a*_*N*_ × *P*_*N*_ − *P*_*MS*_ − *P*_*E*_ represents the total input cost.

#### 2.5.2 Construction and Solution of the Net Profit Target Optimization Model for Crops Grown in Farm X

In the process of constructing the target optimization model for the net profit of crops grown in each plot of Farm X in 2024, some parameters in the model need to be set. For the paddy fields of Farm X, only rice crops are grown, so there is no need to choose which crop will have the highest net profit. However, for the dry fields of Farm X, there are two dry field crops to choose from: maize and soybeans. Therefore, there is a question of which crop will have the highest net profit. In Formula (5-3), let represent the type of crop grown in the i-th dry field of Farm X.

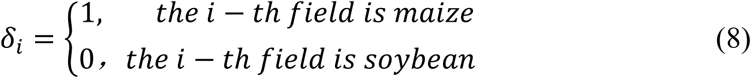

When constructing the optimization model for the net profit target of Farm X in 2024, the objective function was set to maximize the best economic benefits. This means that this study will pursue planting strategies that maximize the net profit of the farm. Net profit refers to the surplus obtained after subtracting the planting cost from the sales revenue of crops. By maximizing net profits, farm managers can ensure that under limited resource conditions, the planting density, fertilizer application amount and other agricultural practices are allocated in the most optimized way, thereby achieving the maximization of the economic benefits of farm production.

For the paddy fields of Farm X, only rice crops are grown, so there is no need to choose which crop will have the highest net profit. However, for the dry fields of Farm X, there are two dry field crops to choose from: maize and soybeans. Therefore, there is a question of which crop will have the highest net profit. Therefore, in formula (9) let represent the net profit of the i-th dry field planted with maize (yuan /ha); represents the net profit of soybean cultivation in the i-th dry field (yuan /ha); represents the area (ha) of the i-th block of Farm X. According to the requirements of Farm X, the total area of maize and soybeans in the farm must not be less than 10,000 hectares. Then the objective function and constraints are:

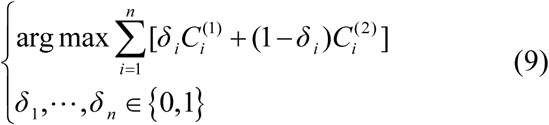

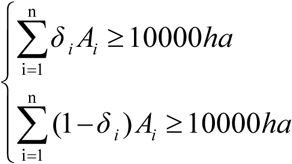

This study developed a planning calculation database that encompasses objective functions and constraint parameters, derived from the planting production structure optimization model based on the net profit target optimization model of X Farm for 2024. The article employs the optimization function of the Genetic Algorithm (GA) to determine the optimal solution for the net profit target optimization model of Farm X in 2024, thereby assessing the net profit associated with crop planting at Farm X. In this research, a genetic algorithm is utilized to identify the most effective planting plan, aimed at optimizing the objective function, which is the net profit of X Farm in 2024.By optimizing the calculations of parameters such as planting density and fertilization amount, the most effective planting strategy can be established in accordance with actual conditions and constraint requirements. The genetic algorithm enhances the characteristics of individuals within the population through a continuous iterative process, thereby progressively converging on the optimal solution. Ultimately, the best planting plan tailored for X farm can be derived to maximize net profit. Following the organization of the calculation results, the net profit values corresponding to each planting strategy can be displayed in the table.

## 3 Results

This study established a planning calculation database including the objective function and constraint condition parameters based on the planting production structure optimization model and the parameters obtained from the net profit target optimization model of Farm X in 2024. This paper utilizes the optimization function of the Genetic Algorithm (GA) to calculate the optimal solution of the net profit target optimization model of Farm X in 2024, to evaluate the net profit of crop planting in Farm X. This study considers the optimal combination of planting density and fertilizer application for the highest net profit under the following three scenarios (free market situation, government subsidy situation, and considering crop rotation and government subsidy situation), thereby determining the optimal planting strategy for Farm X.

In 2024, X Farm comprises 54 distinct plots, each identified by a unique number ranging from 1 to 54. Plots 10 to 21 and 36 to 47 are designated as paddy fields, specifically for rice cultivation. Conversely, plots 1 to 9, 22 to 35, and 48 to 54 are classified as dry land plots, which allow for the cultivation of either maize or soybeans. When determining the optimal planting strategy for the dry land plots of Farm X, it is essential to evaluate the net profits associated with both maize and soybeans.

### 3.1 Planting Strategy Optimization of Farm X under Free Market Conditions

Under the free market, Farm X’s planting strategy for 2024 involves selecting crops based on the principle of maximizing net profit for each plot. Nevertheless, it is essential that the total area allocated to maize and soybeans across all dry fields of Farm X does not fall below 10,000 hectares. By judiciously partitioning the areas designated for paddy and dry fields, Farm X. The strategic allocation of plot characteristics enhances the economic viability of the farm and offers managers greater flexibility in planting decisions and resource management. By optimizing the economic returns from each plot and considering the overall area constraints for maize and soybeans in dryland conditions, Farm X can effectively leverage its land capital to realize its vision of sustained growth while securing economic advantages. The planting strategies for each plot of Farm X in 2024 under the free-market conditions are shown in Table 5.

**Table 5.** Planting strategies for various plots of the Farm X in 2024 under free market conditions.

| Plot Num | Crop type | Fertilizer (kg/ha) | Density(plants/m<sup>2</sup>) | Profit(yuan/ha) |
| --- | --- | --- | --- | --- |
| 1 | Maize | 292.92 | 10.46 | 22768.329 |
| 2 | Maize | 272.46 | 10.50 | 27634.650 |
| 3 | Maize | 338.99 | 9.68 | 18876.469 |
| 4 | Maize | 275.99 | 12.66 | 22232.844 |
| 5 | Maize | 292.48 | 10.11 | 21970.342 |
| 6 | Maize | 337.31 | 8.30 | 19539.254 |
| 7 | Maize | 312.74 | 13.69 | 21257.030 |
| 8 | Maize | 305.15 | 8.42 | 21047.891 |
| 9 | Soybean | 144.33 | 37.18 | 6289.208 |
| 10 | Rice | 411.98 | 46.12 | 9610.390 |
| 11 | Rice | 462.32 | 26.26 | 8623.525 |
| 12 | Rice | 423.44 | 30.17 | 9050.799 |
| 13 | Rice | 348.97 | 45.78 | 9735.240 |
| 14 | Rice | 377.14 | 36.77 | 9845.210 |
| 15 | Rice | 304.66 | 34.09 | 10055.748 |
| 16 | Rice | 355.76 | 42.64 | 9975.302 |
| 17 | Rice | 382.27 | 35.79 | 9686.466 |
| 18 | Rice | 316.05 | 43.31 | 9799.217 |
| 19 | Rice | 317.01 | 38.61 | 9889.923 |
| 20 | Rice | 347.67 | 39.10 | 9911.528 |
| 21 | Rice | 437.24 | 31.23 | 9094.848 |
| 22 | Soybean | 187.82 | 29.64 | 5809.932 |
| 23 | Maize | 269.13 | 11.86 | 21079.480 |
| 24 | Maize | 278.06 | 10.70 | 21573.790 |
| 25 | Maize | 303.22 | 9.36 | 21861.198 |
| 26 | Maize | 366.28 | 9.35 | 18388.833 |
| 27 | Maize | 283.01 | 10.74 | 22513.103 |
| 28 | Soybean | 143.32 | 39.35 | 6219.303 |
| 29 | Soybean | 122.97 | 37.15 | 6335.965 |
| 30 | Soybean | 126.96 | 29.56 | 6412.639 |
| 31 | Maize | 353.90 | 11.19 | 20895.597 |
| 32 | Maize | 352.22 | 14.78 | 19711.456 |
| 33 | Maize | 316.12 | 10.74 | 20905.226 |
| 34 | Maize | 317.73 | 12.63 | 20381.261 |
| 35 | Soybean | 161.00 | 28.60 | 6149.899 |
| 36 | Rice | 402.45 | 45.91 | 9451.460 |
| 37 | Rice | 342.26 | 41.26 | 10049.804 |
| 38 | Rice | 316.99 | 38.20 | 10078.505 |
| 39 | Rice | 306.94 | 49.77 | 9601.216 |
| 40 | Rice | 351.34 | 43.61 | 9729.832 |
| 41 | Rice | 318.34 | 43.42 | 9852.498 |
| 42 | Rice | 366.46 | 41.89 | 9621.320 |
| 43 | Rice | 397.26 | 36.62 | 9742.677 |
| 44 | Rice | 310.48 | 45.95 | 10893.872 |
| 45 | Rice | 314.63 | 38.77 | 10051.063 |
| 46 | Rice | 460.36 | 34.57 | 8866.663 |
| 47 | Rice | 320.20 | 39.40 | 9889.379 |
| 48 | Soybean | 179.63 | 38.83 | 5807.037 |
| 49 | Soybean | 174.00 | 33.73 | 5786.247 |
| 50 | Maize | 264.08 | 9.23 | 20021.424 |
| 51 | Soybean | 121.92 | 38.48 | 6000.867 |
| 52 | Maize | 265.28 | 11.19 | 20189.534 |
| 53 | Soybean | 175.98 | 38.79 | 5847.788 |
| 54 | Maize | 315.81 | 10.19 | 20762.746 |

The maize variety grown in Farm X is Demeiya 3, the rice variety is Longjing 1624, and the soybean variety is Kendou 94. The total net profit (in yuan) of crop cultivation in each plot of Farm X in 2024 under the free market conditions is shown in Figure 8.

**Figure 8.**
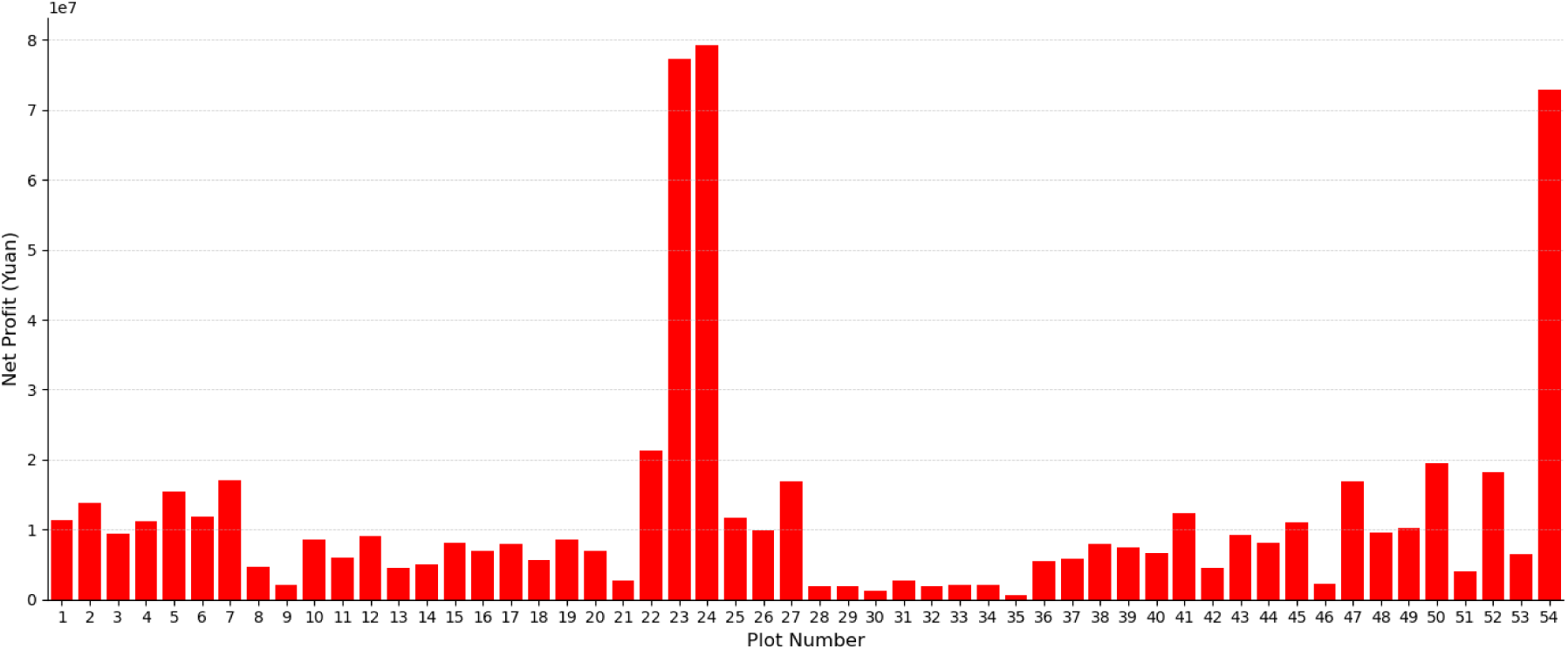
Under free market conditions, the total net profit of crop cultivation on each plot of the Farm X in 2024 (yuan)

Utilizing the mentioned planting data for each plot within the free market environment, a detailed crop planting strategy for X Farm in 2024 can be devised. This strategy encompasses specific arrangements for crop planting density and fertilizer application across each plot, which will be represented in practical and detailed prescription maps. These maps will enable Farm X to optimize automated mechanical sowing, thereby facilitating large-scale agricultural production. The prescription chart illustrating crop planting density (plants/m²) for each plot at Farm X in 2024 within the free-market context is presented in Figure 3-2. Additionally, the prescription diagram detailing the amount of fertilizer (kg/ha) required for each plot at Farm X in 2024 under the free market is also depicted in Figure 9.

**Figure 9.**
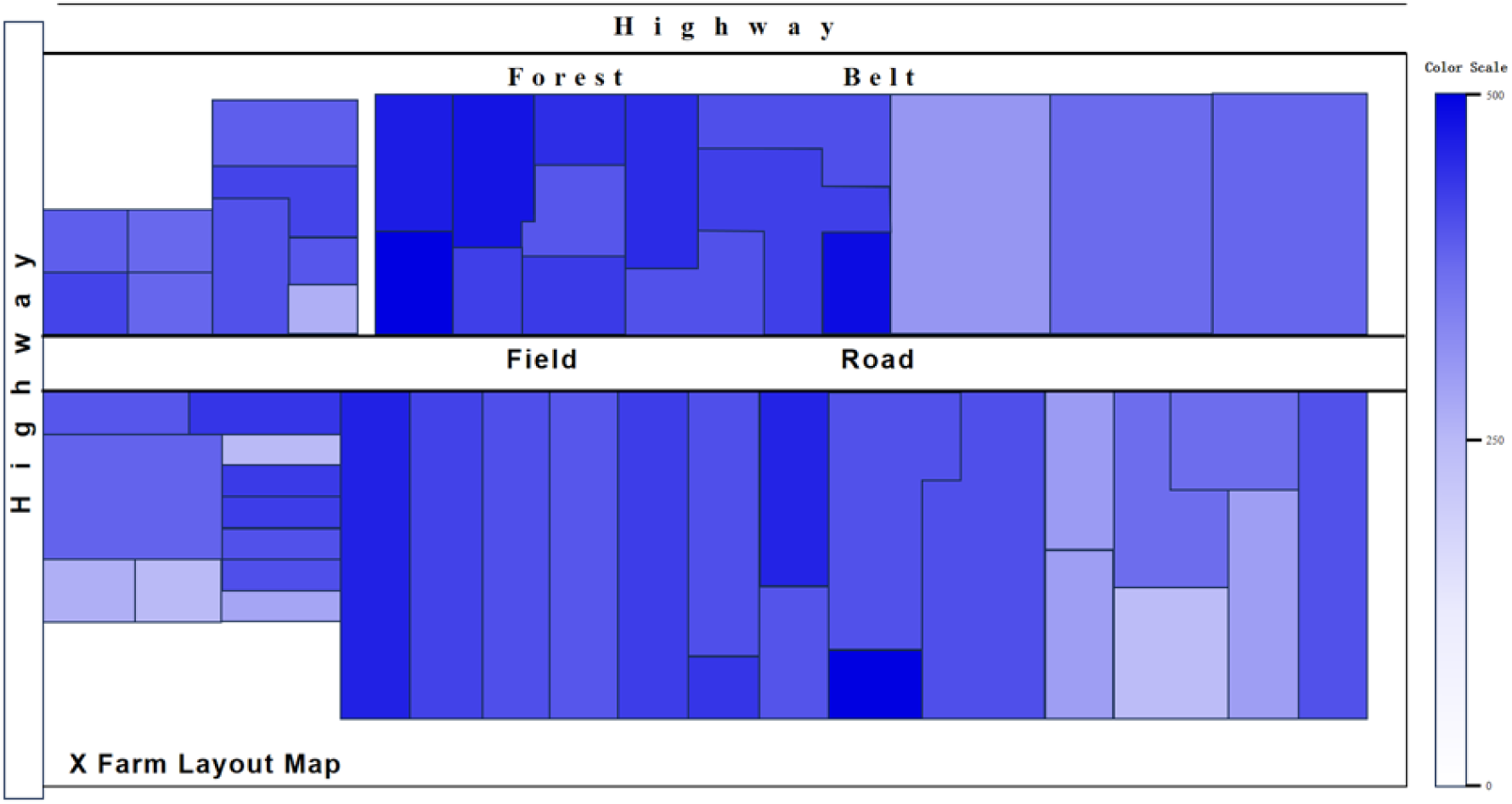
Prescription chart of crop planting density for each plot of the Farm X in 2024 under free market conditions (plants/m^2^)

### 3.2 Planting Strategy Optimization of the dryland plots of Farm X

Planting Strategy of the dryland plots of Farm X, it is essential to evaluate the net profit of maize and soybeans to determine the most appropriate crop planting strategy. As previously noted, plots 10 to 21 and 36 to 47 are allocated for rice cultivation, whereas plots 1 to 9, 22 to 35, and 48 to 54 are designated as dryland areas suitable for the cultivation of either maize or soybeans. In the event of government subsidies, the planting strategy of Farm. It is essential to ensure that the total area allocated to maize and soybeans does not fall below 10,000 hectares. In 2024, maximizing the economic potential of each plot on Farm X must consider the total area limitations for maize and soybeans in dryland conditions. The formulation of the planting strategy for Farm X in 2024, in conjunction with government subsidies, is guided by the "Notice of the Heilongjiang Provincial Department of Finance on Allocating Subsidy Funds to Maize, Soybean and Rice Producers." According to this notice, the subsidy rate for maize producers is set at 14 yuan per mu, while soybean producers receive a subsidy of 366 yuan per mu, and rice surface water irrigation is subsidized at 172.67 yuan per mu. The planting strategy for each plot of Farm X in 2024, incorporating these government subsidies, is detailed in Table 6.

**Table 6.** Planting strategies for various plots of the Farm X in 2024 under government subsidies.

| Plot Num | Crop Type | Fertilizer (kg/ha) | Density(plants/m<sup>2</sup>) | Profit(yuan/ha) |
| --- | --- | --- | --- | --- |
| 1 | Soybean | 189.60 | 31.78 | 11484.064 |
| 2 | Maize | 331.65 | 9.97 | 26696.890 |
| 3 | Soybean | 167.78 | 26.82 | 11581.767 |
| 4 | Soybean | 199.40 | 29.30 | 11856.030 |
| 5 | Maize | 327.25 | 10.91 | 21839.545 |
| 6 | Maize | 331.37 | 13.72 | 20063.748 |
| 7 | Maize | 282.22 | 11.45 | 22248.171 |
| 8 | Soybean | 158.40 | 27.07 | 11735.144 |
| 9 | Maize | 331.09 | 13.09 | 21962.652 |
| 10 | Rice | 338.22 | 36.84 | 12668.184 |
| 11 | Rice | 329.69 | 47.34 | 12317.216 |
| 12 | Rice | 359.24 | 39.30 | 12318.428 |
| 13 | Rice | 489.04 | 43.41 | 11547.647 |
| 14 | Rice | 372.54 | 37.99 | 12466.908 |
| 15 | Rice | 313.48 | 48.62 | 12326.011 |
| 16 | Rice | 413.30 | 44.47 | 12249.408 |
| 17 | Rice | 328.98 | 41.59 | 12522.122 |
| 18 | Rice | 327.20 | 28.52 | 12085.330 |
| 19 | Rice | 410.13 | 32.50 | 11824.796 |
| 20 | Rice | 333.26 | 37.86 | 12553.327 |
| 21 | Rice | 371.71 | 45.14 | 12181.302 |
| 22 | Soybean | 122.11 | 36.95 | 11612.382 |
| 23 | Maize | 281.49 | 11.70 | 21368.219 |
| 24 | Maize | 251.48 | 10.93 | 21657.461 |
| 25 | Soybean | 181.99 | 25.48 | 11592.056 |
| 26 | Soybean | 145.92 | 29.99 | 11772.080 |
| 27 | Maize | 274.39 | 12.41 | 22607.834 |
| 28 | Soybean | 128.51 | 27.74 | 11809.956 |
| 29 | Soybean | 169.70 | 35.47 | 11600.926 |
| 30 | Maize | 319.99 | 14.38 | 20711.461 |
| 31 | Soybean | 130.27 | 32.55 | 11871.966 |
| 32 | Maize | 295.96 | 10.46 | 22028.751 |
| 33 | Soybean | 176.30 | 26.25 | 11224.781 |
| 34 | Maize | 377.02 | 11.70 | 20440.298 |
| 35 | Soybean | 188.25 | 30.76 | 11451.946 |
| 36 | Rice | 380.26 | 32.99 | 12220.467 |
| 37 | Rice | 360.35 | 38.68 | 12601.979 |
| 38 | Rice | 375.83 | 47.64 | 12172.732 |
| 39 | Rice | 371.52 | 36.93 | 12418.911 |
| 40 | Rice | 462.97 | 43.56 | 11638.590 |
| 41 | Rice | 332.81 | 39.57 | 12466.342 |
| 42 | Rice | 322.25 | 34.63 | 12418.680 |
| 43 | Rice | 325.70 | 45.28 | 12051.305 |
| 44 | Rice | 302.57 | 42.05 | 13537.090 |
| 45 | Rice | 316.21 | 38.63 | 12637.111 |
| 46 | Rice | 436.73 | 42.93 | 11735.240 |
| 47 | Rice | 362.26 | 42.72 | 12225.803 |
| 48 | Soybean | 195.77 | 35.83 | 11243.723 |
| 49 | Maize | 373.86 | 11.34 | 20783.864 |
| 50 | Soybean | 197.20 | 34.86 | 11256.131 |
| 51 | Maize | 365.10 | 9.18 | 19778.543 |
| 52 | Maize | 354.19 | 11.26 | 21096.781 |
| 53 | Maize | 259.90 | 9.51 | 20329.442 |
| 54 | Maize | 321.98 | 11.63 | 21028.149 |

The maize variety planted on Farm X is **Demeiya No. 3**, the rice variety is **Longjing 1624**, and the soybean variety is **Kendou 94**. **Figure 11** shows the total net profit (in yuan) of crop planting on each plot of Farm X in 2024 under government subsidies.

**Figure 10.**
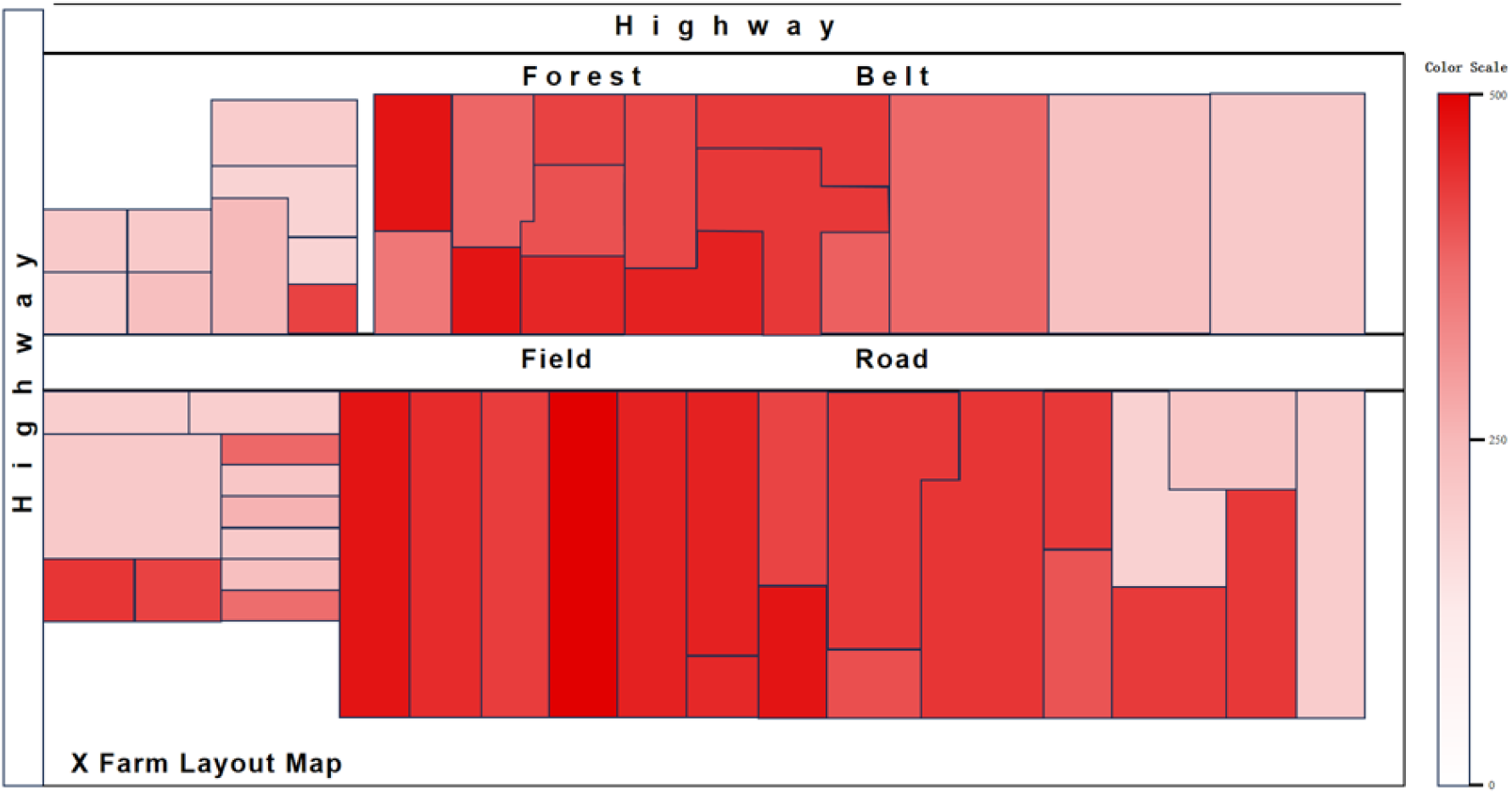
Prescription chart of crop fertilization amount for each plot of the Farm X in 2024 under free market conditions (kg/ha)

**Figure 11.**
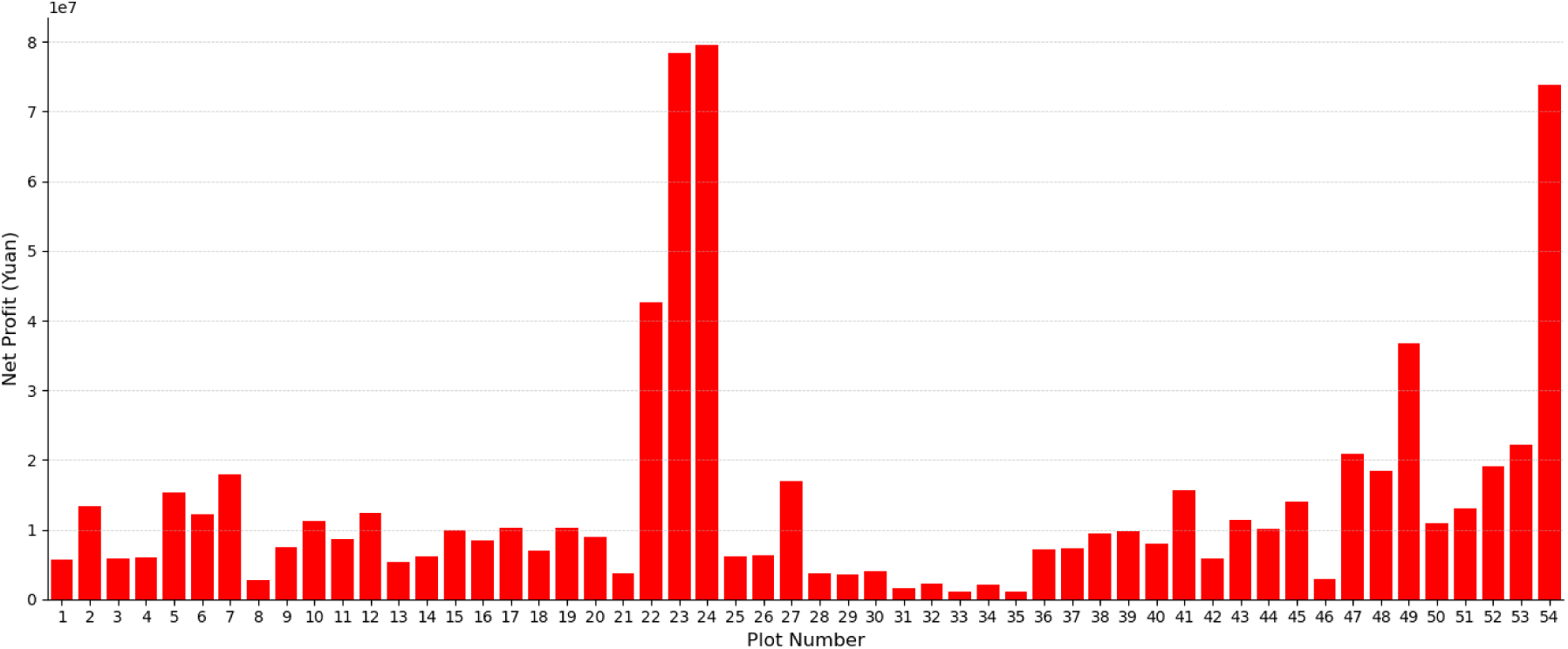
Under government subsidies, the total net profit of crop cultivation on each plot of the Farm X in 2024 (yuan)

Based on the government subsidy information obtained above and the planting data of each plot, a detailed crop planting strategy and prescription map can be formulated for Farm X in 2024 under the advantage of government subsidies. This strategy will cover the crop planting density and fertilizer application amount of each plot and fully utilize the government subsidy policy, which can achieve efficient operation of farm production. The prescription diagram of crop planting density (plants/m2) for each plot of Farm X in 2024 under government subsidies is shown in Figure 3-6. The prescription chart of crop fertilizer application (kg/ha) for each plot of Farm X in 2024 under government subsidies is shown in Figure 12.

**Figure 12.**
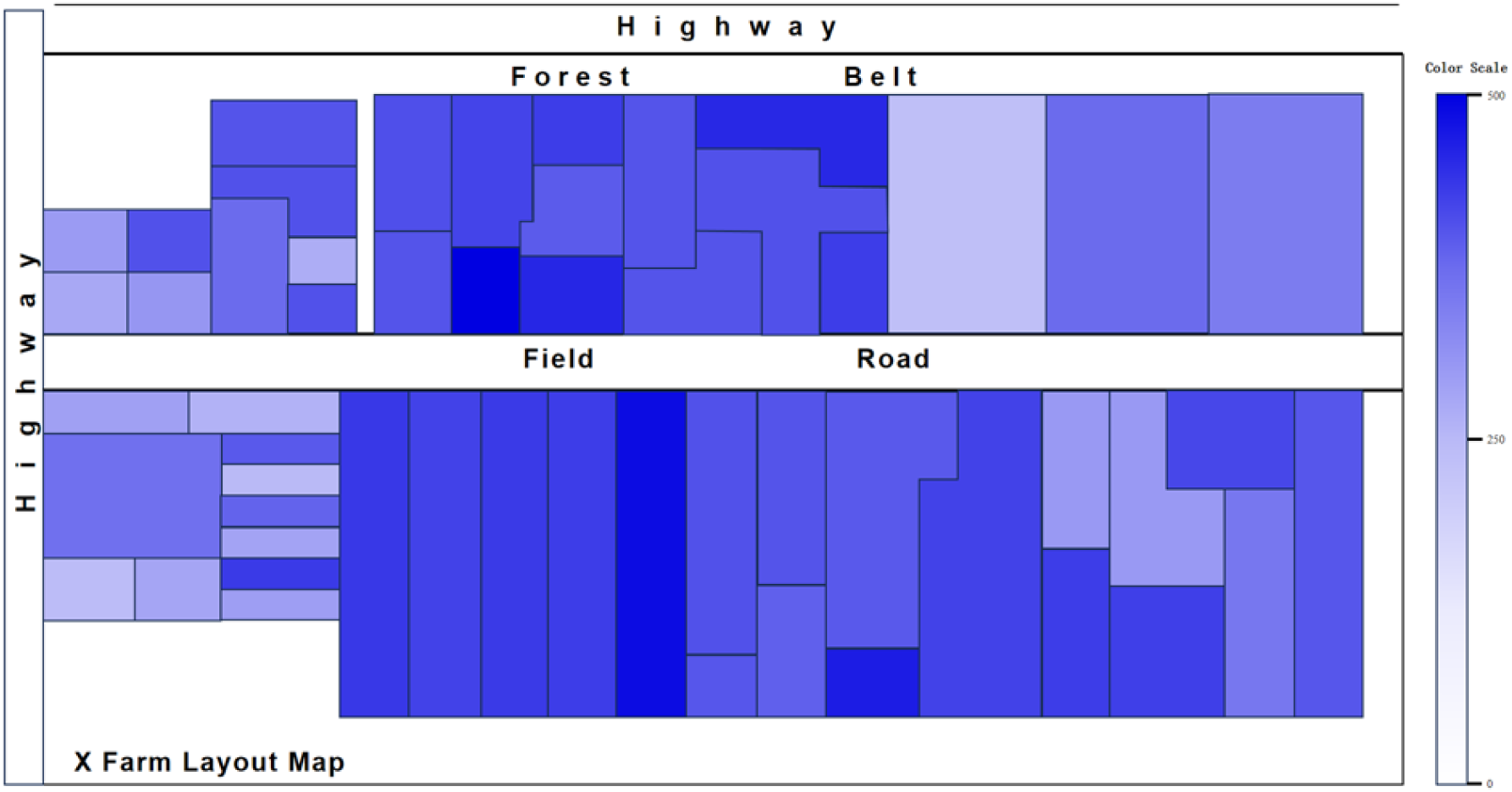
Prescription chart of crop planting density for each plot of the Farm X in 2024 under government subsidies (plants/m^2^)

### 3.3 Consider the planting strategies of Farm X under crop rotation and government subsidies

When devising the planting strategy for Farm X, it is essential to establish an effective crop rotation. The selection of crops for each plot in 2024 must be informed by the types of crops cultivated in 2023. Notably, the crops in the paddy fields will remain unchanged, continuing to be rice in 2024. For dry land, appropriate rotation crops should be chosen based on the crop type from the previous year.

Through crop rotation, farms can enhance soil fertility, increase crop yields, and mitigate pest and disease issues. This strategic allocation of plot characteristics not only improves farm productivity and profitability but also offers farm managers greater flexibility in planting decisions and resource management. By optimizing the economic benefits of each plot, Farm X can fully utilize its land resources and attain sustainable development alongside economic gains. This section of the study must consider both crop rotation and government subsidies. The planting strategy for each plot of Farm X in 2024, considering these factors, is presented in Table 7.

**Table 7.**
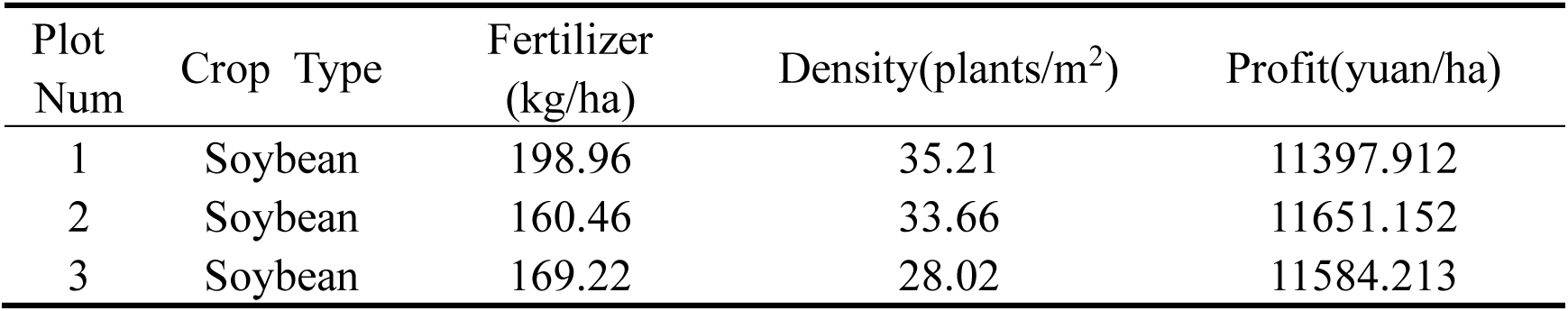
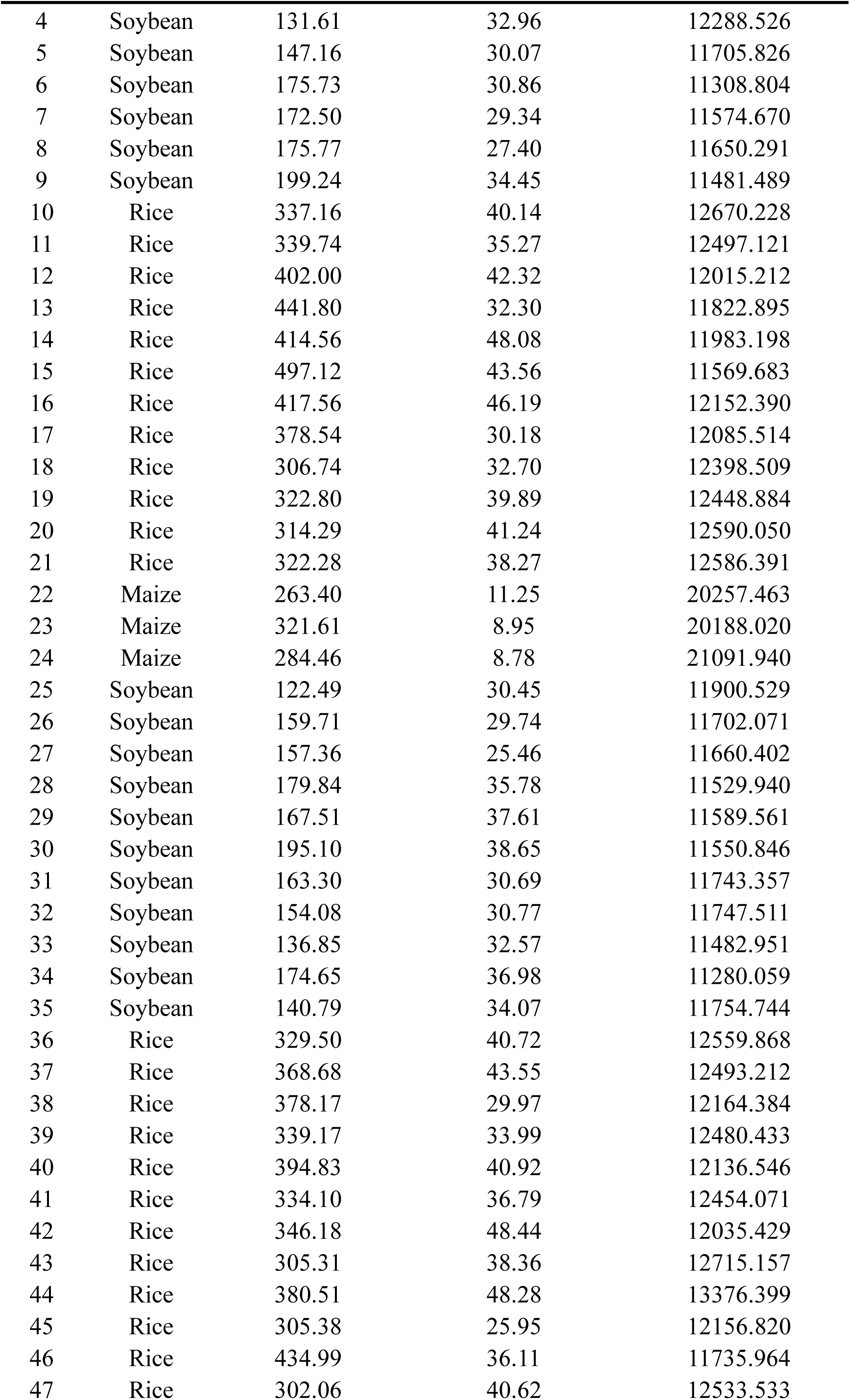
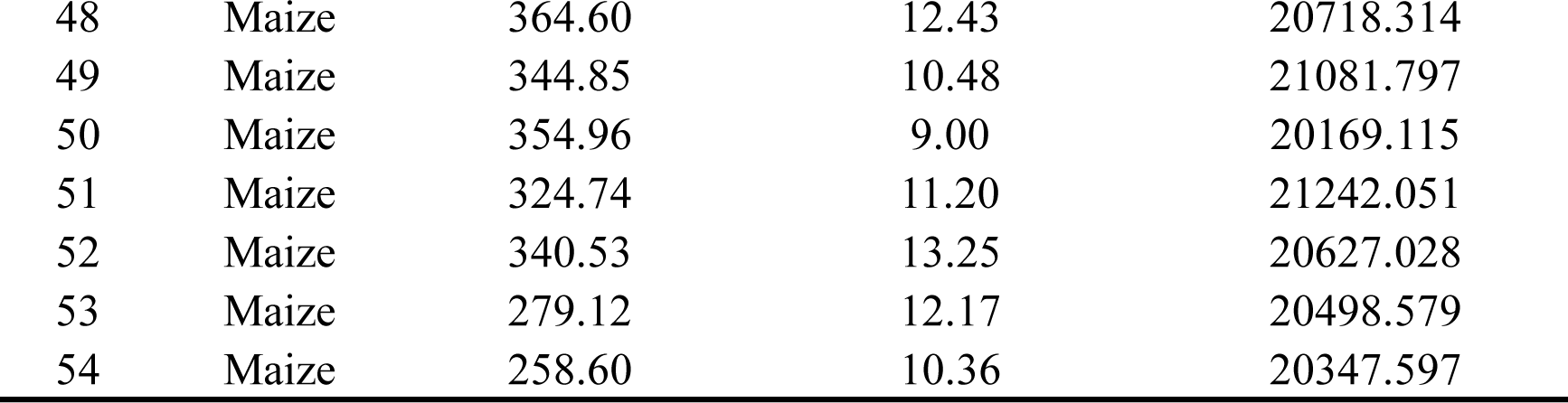
Considering crop rotation and government subsidies, planting strategies for each plot of the Farm X in 2024.

| Plot Num | Crop Type | Fertilizer (kg/ha) | Density(plants/m<sup>2</sup>) | Profit(yuan/ha) |
| --- | --- | --- | --- | --- |
| 1 | Soybean | 198.96 | 35.21 | 11397.912 |
| 2 | Soybean | 160.46 | 33.66 | 11651.152 |
| 3 | Soybean | 169.22 | 28.02 | 11584.213 |
| 4 | Soybean | 131.61 | 32.96 | 12288.526 |
| 5 | Soybean | 147.16 | 30.07 | 11705.826 |
| 6 | Soybean | 175.73 | 30.86 | 11308.804 |
| 7 | Soybean | 172.50 | 29.34 | 11574.670 |
| 8 | Soybean | 175.77 | 27.40 | 11650.291 |
| 9 | Soybean | 199.24 | 34.45 | 11481.489 |
| 10 | Rice | 337.16 | 40.14 | 12670.228 |
| 11 | Rice | 339.74 | 35.27 | 12497.121 |
| 12 | Rice | 402.00 | 42.32 | 12015.212 |
| 13 | Rice | 441.80 | 32.30 | 11822.895 |
| 14 | Rice | 414.56 | 48.08 | 11983.198 |
| 15 | Rice | 497.12 | 43.56 | 11569.683 |
| 16 | Rice | 417.56 | 46.19 | 12152.390 |
| 17 | Rice | 378.54 | 30.18 | 12085.514 |
| 18 | Rice | 306.74 | 32.70 | 12398.509 |
| 19 | Rice | 322.80 | 39.89 | 12448.884 |
| 20 | Rice | 314.29 | 41.24 | 12590.050 |
| 21 | Rice | 322.28 | 38.27 | 12586.391 |
| 22 | Maize | 263.40 | 11.25 | 20257.463 |
| 23 | Maize | 321.61 | 8.95 | 20188.020 |
| 24 | Maize | 284.46 | 8.78 | 21091.940 |
| 25 | Soybean | 122.49 | 30.45 | 11900.529 |
| 26 | Soybean | 159.71 | 29.74 | 11702.071 |
| 27 | Soybean | 157.36 | 25.46 | 11660.402 |
| 28 | Soybean | 179.84 | 35.78 | 11529.940 |
| 29 | Soybean | 167.51 | 37.61 | 11589.561 |
| 30 | Soybean | 195.10 | 38.65 | 11550.846 |
| 31 | Soybean | 163.30 | 30.69 | 11743.357 |
| 32 | Soybean | 154.08 | 30.77 | 11747.511 |
| 33 | Soybean | 136.85 | 32.57 | 11482.951 |
| 34 | Soybean | 174.65 | 36.98 | 11280.059 |
| 35 | Soybean | 140.79 | 34.07 | 11754.744 |
| 36 | Rice | 329.50 | 40.72 | 12559.868 |
| 37 | Rice | 368.68 | 43.55 | 12493.212 |
| 38 | Rice | 378.17 | 29.97 | 12164.384 |
| 39 | Rice | 339.17 | 33.99 | 12480.433 |
| 40 | Rice | 394.83 | 40.92 | 12136.546 |
| 41 | Rice | 334.10 | 36.79 | 12454.071 |
| 42 | Rice | 346.18 | 48.44 | 12035.429 |
| 43 | Rice | 305.31 | 38.36 | 12715.157 |
| 44 | Rice | 380.51 | 48.28 | 13376.399 |
| 45 | Rice | 305.38 | 25.95 | 12156.820 |
| 46 | Rice | 434.99 | 36.11 | 11735.964 |
| 47 | Rice | 302.06 | 40.62 | 12533.533 |
| 48 | Maize | 364.60 | 12.43 | 20718.314 |
| 49 | Maize | 344.85 | 10.48 | 21081.797 |
| 50 | Maize | 354.96 | 9.00 | 20169.115 |
| 51 | Maize | 324.74 | 11.20 | 21242.051 |
| 52 | Maize | 340.53 | 13.25 | 20627.028 |
| 53 | Maize | 279.12 | 12.17 | 20498.579 |
| 54 | Maize | 258.60 | 10.36 | 20347.597 |

Based on the planting data of each plot obtained above, considering crop rotation and government subsidies, a detailed crop planting strategy prescription map can be formulated for Farm X in 2024 under the advantages of crop rotation and government subsidies. This strategy prescription map will cover the crop planting density and fertilizer application amount of each plot in Farm X, so as to fully utilize the government subsidy policy and achieve efficient operation of agricultural production. The total net profit (yuan) of crop cultivation in each plot of Farm X in 2024, considering crop rotation and government subsidies, is shown in Figure 14.

**Figure 13.**
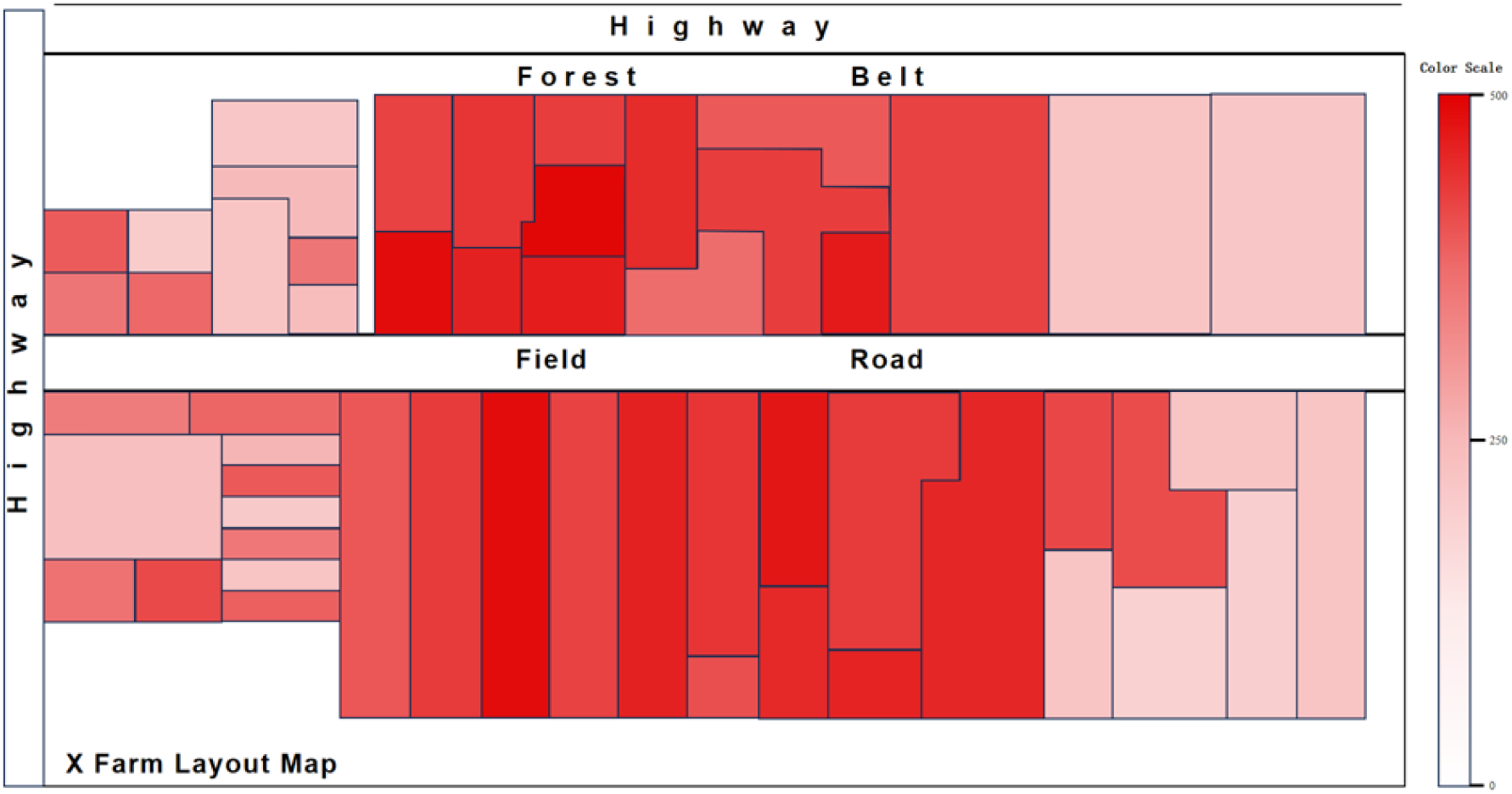
Prescription chart of crop fertilization amount for each plot of the Farm X in 2024 under government subsidies (kg/ha)

**Figure 14.**
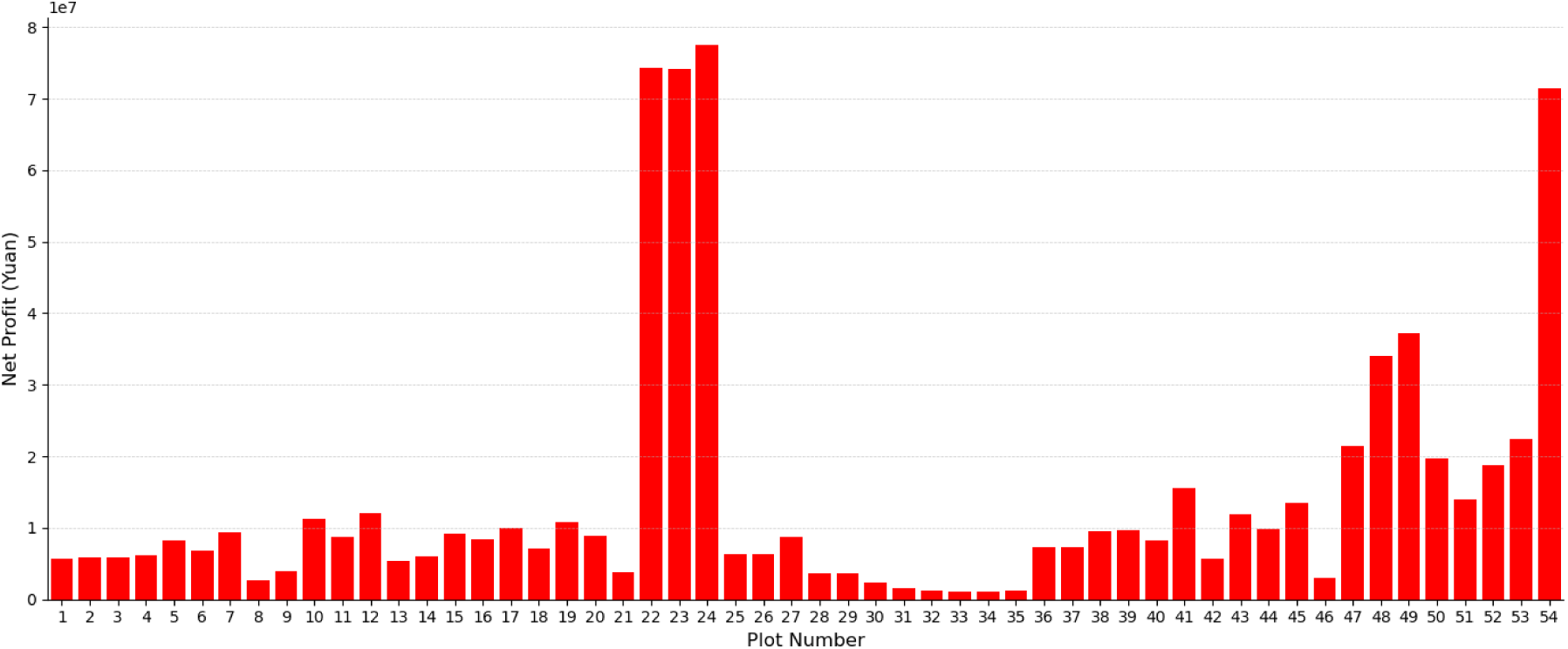
Total Net Profit of Each Plot on Farm X in 2024 under Crop Rotation and Government Subsidies (yuan)

The prescription diagram of crop planting density (plants/m2) for each plot of Farm X in 2024 under government subsidies is shown in Figure 15. The prescription chart of crop fertilizer application (kg/ha) for each plot of Farm X in 2024 under government subsidies is shown in Figure 16.

**Figure 15.**
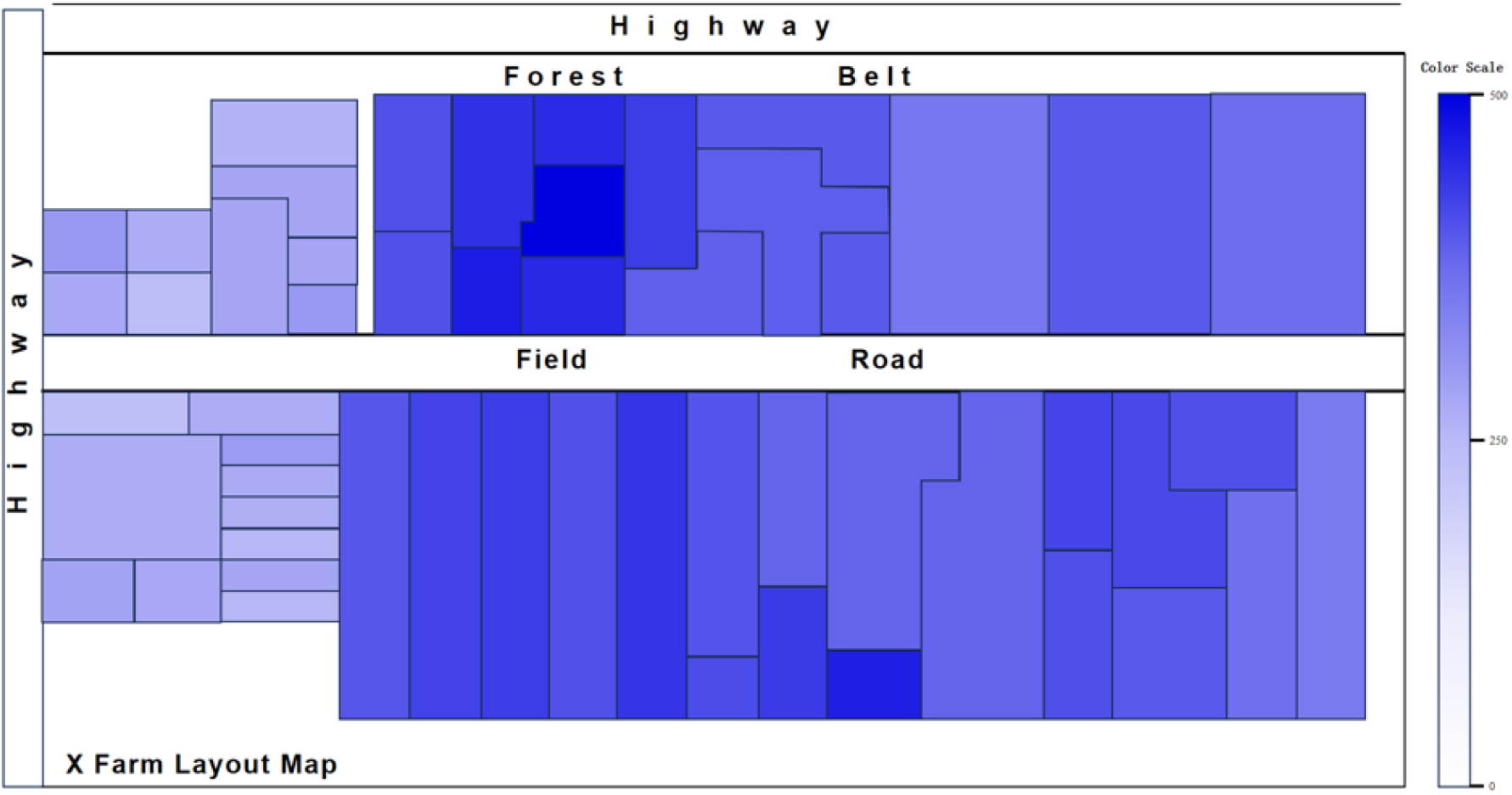
Planting Density Prescription for Each Plot of Farm X in 2024 under Crop Rotation and Government Subsidies (kg/ha)

**Figure 16.**
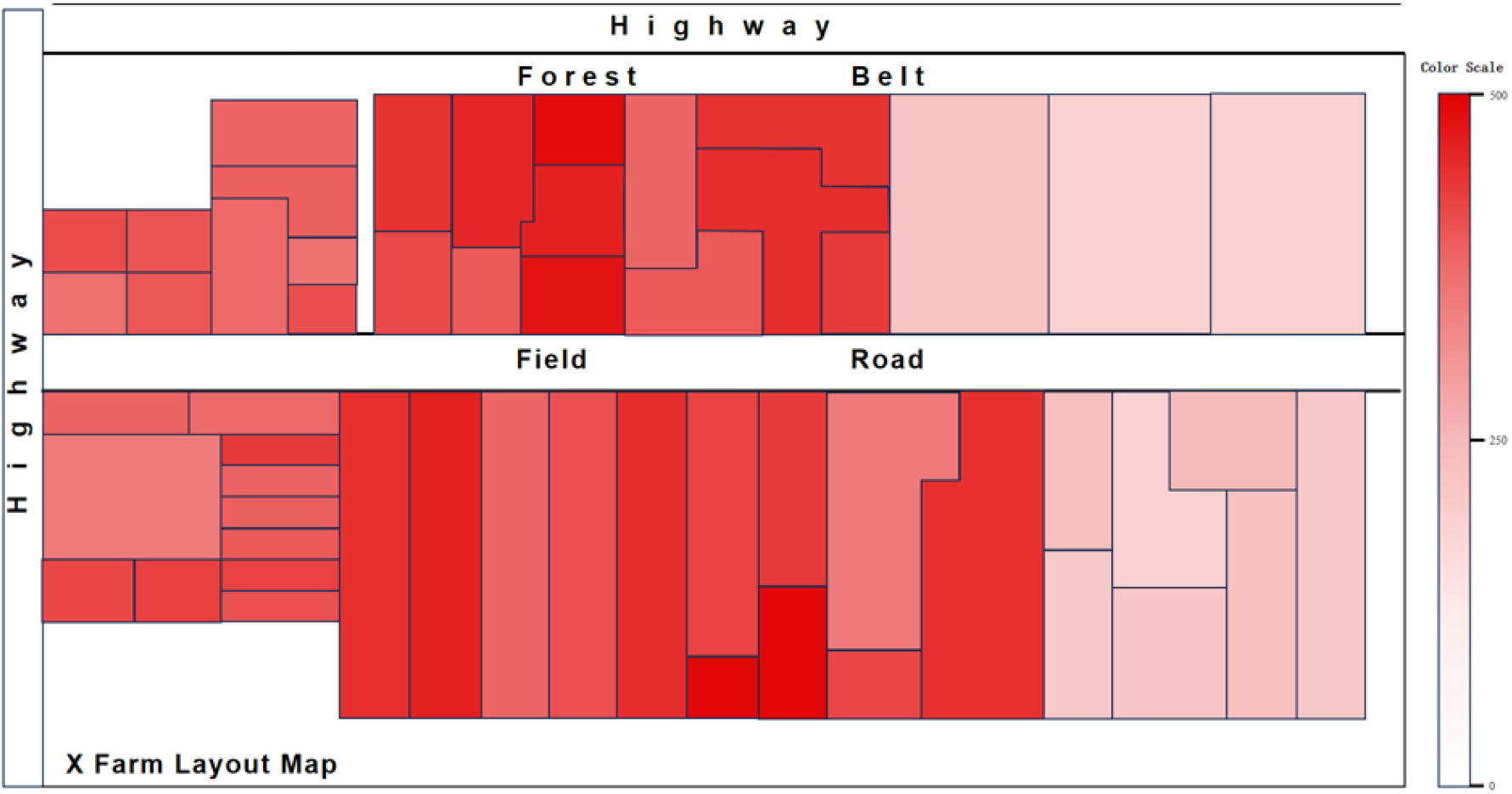
Fertilizer Prescription for Each Plot of Farm X in 2024 under Crop Rotation and Government Subsidies (kg/ha)

## 4 Discussion

This study presents a robust method for optimizing the intelligent planting strategy of X farm. Future research may be extended to encompass areas such as data updating, multi-model fusion, market price fluctuations, and policy factors, thereby enhancing the overall efficiency and quality of agricultural production in our country. Concurrently, the research findings can be integrated with intelligent decision support systems to facilitate more efficient and intelligent agricultural management.

This study undertook a systematic investigation of the planting strategy employed by X farm, utilizing the DSSAT model to optimize and analyze the planting strategy for 2023, while also providing a detailed prediction and optimization for 2024. The research offers effective optimization solutions for farms, thereby enhancing the efficiency of agricultural production and resource utilization. Nonetheless, despite the valuable insights presented, certain limitations and opportunities for further development warrant attention and should be addressed in subsequent research. Primarily, the data source underpins this study, with the information derived from the experimental data of X farm. As farm and crop conditions evolve over time, the data may also be subject to change. To ensure the timeliness and accuracy of planting strategies, subsequent research must place greater emphasis on data updates. In large-scale agricultural production, factors such as seasonal variations, soil conditions, and climate change can significantly impact data. Therefore, future studies should continuously monitor and update data to enable real-time adjustments to models, thereby facilitating more precise optimization of planting strategies. Furthermore, although this study employed the DSSAT model for the optimization analysis of planting strategies, numerous other advanced planting models are currently available on the market. For instance, the Agricultural Production Systems Simulator (APSIM) and the STICS model have each demonstrated strong adaptability and predictive capabilities under varying environmental conditions. Consequently, the integration or comparison of multiple implant models can offer diverse perspectives and validations for research, thereby augmenting the applicability of the DSSAT model. The fusion of various models can also enhance adaptability to the specific conditions of different farms, further improving the precision of planting strategies. Moreover, the impact of market price fluctuations and policy factors on planting decisions warrants attention. Fluctuations in crop prices directly influence farmers’ planting choices, while national or regional agricultural policies and subsidy schemes also shape the selection of planting strategies. Therefore, future research should aim to incorporate market price fluctuations and policy factors into the model to render the planting strategy more comprehensive and applicable. By taking these external factors into account, the model can enhance the accuracy of predictions when optimizing planting strategies, thereby assisting farmers in making more advantageous decisions in response to market fluctuations and policy modifications. Furthermore, the scope of this study was confined to Farm X and primarily examined several crops, including rice, maize, and soybeans. While these crops are representative of Farm X, they encompass only a fraction of agricultural production. Future research could extend the findings of this study to other farms and crops, particularly those situated in diverse climatic zones and soil conditions across the country. By broadening the range of agricultural environments considered, the applicability and generalizability of the research may be enhanced, ultimately contributing to the overall efficiency and quality of agricultural production within the country. Finally, while this study offers farms optimization options for planting strategies, an intelligent decision support system has yet to be developed. With advancements in information technology and artificial intelligence, agricultural production is progressively entering an era of intelligence. Future research should aim to integrate the findings of this study with intelligent decision support systems to create a platform grounded in data analysis and model optimization. Such a system would be capable of collecting and processing diverse data types in real time, providing optimization recommendations to farm managers through model predictions, thus facilitating more efficient and intelligent agricultural management.

This study presents an effective approach for optimizing the intelligent planting strategy of X farm and delineates a pathway for future research. The incorporation of data updates, multi-model fusion, market price fluctuations, and policy factors is anticipated to enhance the accuracy and adaptability of planting strategies in subsequent investigations. Concurrently, the advancement of intelligent decision support systems may facilitate more efficient and intelligent agricultural management, thereby offering crucial support for the modernization and sustainable development of agriculture in our country.

## 5 Conclusion

The study utilized X Farm in Heilongjiang Province as a case study, concentrating on the simulation and optimization of the intelligent planting strategy through the DSSAT model, with the objective of achieving dual optimization of economic benefits and resource efficiency. The DSSAT model was employed to optimize and analyze the planting strategy for X Farm in 2023. To ensure comprehensive experimental data from each plot at X Farm, including meteorological data, soil characteristics, crop genetic parameters, and management practices, this study devised a systematic experimental plan. This plan encompassed six distinct fertilization treatments and six varying planting densities, resulting in a total of 36 detailed planting schemes. Utilizing the DSSAT model, this study simulates the yields of rice, maize, and soybean crops on the Farm. Specifically, for plots cultivated with maize, the optimal solution involves a planting density of approximately 12 (plants/m²) and a fertilizer application of around 300-400 (kg/ha). Notably, plot 29 achieves the highest net profit per unit, amounting to 14,783 (yuan/ha). In the case of rice-growing plots, the ideal solution comprises a planting density of about 25-30 (plants/m²) and a fertilizer dosage of approximately 400 (kg/ha). Among these, plot 15 records the highest net profit per unit, reaching 11,319 (yuan/ha).For plots cultivated with soybeans, the optimal planting density is approximately 35-40 (plants/m²) alongside a fertilizer application of around 320 (kg/ha). Notably, plot No. 51 exhibits the highest net profit per unit, amounting to 9359 (yuan/ha). Utilizing the DSSAT model, predictions were made regarding the unit yield of crops planted across various plots of farm X in 2024, facilitating a refined optimization analysis of the farm’s planting strategy. Chapter 4 of this article continues to employ the 36 meticulously differentiated crop planting plans previously constructed, which encompass various fertilization treatments and interactions of differing planting densities. Subsequently, statistical analysis was performed using SPSS, allowing for the exploration of the internal relationship model between the unit yield of crops planted in diverse plots of farm X in 2024, planting density, and fertilization amounts, derived from the collected data.。 Utilizing these models, the principal objective is to maximize the unit yield of each plot within the Farm.

This study aims to construct a model that elucidates the relationship between crop unit yield, planting density, and fertilization amount for each plot of farmland. To achieve this, comprehensive data on market prices and planting costs of food crops in Heilongjiang Province over the years were collected. Subsequently, SPSS was employed to conduct an economic analysis and to forecast the price and planting costs of food crops in Heilongjiang Province for 2024. Using the cost-benefit analysis method, the study examined the relationship model between the net profit per unit of crops in each plot, planting density, and fertilization amount under various scenarios, including a free market context, a government subsidy scenario, and a situation that accounts for both crop rotation and government subsidies. Following the application of genetic algorithms (GA) to address these models, we summarized the optimal planting strategies that yield the highest net profit for each plot across various scenarios, including free market conditions, government subsidy situations, and considerations of crop rotation alongside government subsidies. We subsequently converted these strategies into intuitive prescription maps that detail optimal planting densities and fertilizer amounts. These maps serve as a foundation for farm machinery and automated farming, while also offering practical decision support tools for farmers. Such tools enable the formulation of scientifically informed planting management strategies that adapt to market fluctuations, ultimately facilitating the dual optimization of economic benefits and resource efficiency.

## Appendix 1

**S-Table 1.** Regression model for unit yield of maize crops planted in 290 Farm in 2024.

| N | Regression Model | R<sup>2</sup> | F value | P value |
| --- | --- | --- | --- | --- |
| 1 | $Y = -0.080X_1^2 + 32.207X_1 - 65.612X_2^2 + 1159.199X_2 + 1.405X_1X_2 - 334.780$ | 0.941 | 215.274 | <0.01 |
| 2 | $Y = -0.081X_1^2 + 32.192X_1 - 69.469X_2^2 + 1290.861X_2 + 1.383X_1X_2 - 1343.827$ | 0.946 | 104.608 | <0.01 |
| 3 | $Y = -0.053X_1^2 + 22.229X_1 - 62.091X_2^2 + 1303.812X_2 + 0.711X_1X_2 + 201.831$ | 0.952 | 119.815 | <0.01 |
| 4 | $Y = -0.082X_1^2 + 37.375X_1 - 56.021X_2^2 + 1025.351X_2 + 0.833X_1X_2 + 932.880$ | 0.936 | 88.094 | <0.01 |
| 5 | $Y = -0.087X_1^2 + 38.745X_1 - 60.471X_2^2 + 1111.648X_2 + 0.985X_1X_2 - 8.387$ | 0.937 | 89.125 | <0.01 |
| 6 | $Y = -0.074X_1^2 + 37.721X_1 - 51.364X_2^2 + 791.318X_2 + 1.075X_1X_2 + 688.604$ | 0.954 | 125.080 | <0.01 |
| 7 | $Y = -0.093X_1^2 + 43.503X_1 - 55.078X_2^2 + 872.217X_2 + 1.221X_1X_2 + 273.110$ | 0.951 | 116.362 | <0.01 |
| 8 | $Y = -0.089X_1^2 + 39.696X_1 - 57.254X_2^2 + 1005.730X_2 + 0.996X_1X_2 + 735.191$ | 0.930 | 79.794 | <0.01 |
| 9 | $Y = -0.099X_1^2 + 44.849X_1 - 56.703X_2^2 + 867.787X_2 + 1.363X_1X_2 + 308.002$ | 0.942 | 97.215 | <0.01 |
| 22 | $Y = -0.080X_1^2 + 35.510X_1 - 54.274X_2^2 + 877.840X_2 + 1.116X_1X_2 + 942.128$ | 0.929 | 78.345 | <0.01 |
| 23 | $Y = -0.101X_1^2 + 47.225X_1 - 52.576X_2^2 + 864.996X_2 + 0.999X_1X_2 + 15.901$ | 0.931 | 80.995 | <0.01 |
| 24 | $Y = -0.103X_1^2 + 48.135X_1 - 53.128X_2^2 + 891.084X_2 + 0.964X_1X_2 - 71.350$ | 0.924 | 73.145 | <0.01 |
| 25 | $Y = -0.080X_1^2 + 36.425X_1 - 56.318X_2^2 + 1038.631X_2 + 0.817X_1X_2 + 1037.921$ | 0.933 | 83.892 | <0.01 |
| 26 | $Y = -0.064X_1^2 + 27.874X_1 - 53.696X_2^2 + 1038.866X_2 + 0.564X_1X_2 + 2085.338$ | 0.913 | 63.112 | <0.01 |
| 27 | $Y = -0.079X_1^2 + 35.008X_1 - 57.653X_2^2 + 1070.367X_2 + 0.822X_1X_2 + 1111.581$ | 0.928 | 77.189 | <0.01 |
| 28 | $Y = -0.077X_1^2 + 34.900X_1 - 56.450X_2^2 + 1054.721X_2 + 0.773X_1X_2 + 1169.016$ | 0.933 | 84.182 | <0.01 |
| 29 | $Y = -0.067X_1^2 + 29.927X_1 - 58.179X_2^2 + 1117.135X_2 + 0.683X_1X_2 + 1596.132$ | 0.934 | 84.278 | <0.01 |
| 30 | $Y = -0.093X_1^2 + 41.762X_1 - 58.292X_2^2 + 933.349X_2 + 1.236X_1X_2 + 462.545$ | 0.940 | 93.549 | <0.01 |
| 31 | $Y = -0.091X_1^2 + 40.310X_1 - 59.825X_2^2 + 971.498X_2 + 1.207X_1X_2 + 553.851$ | 0.936 | 87.131 | <0.01 |
| 32 | $Y = -0.094X_1^2 + 42.188X_1 - 58.799X_2^2 + 888.952X_2 + 1.399X_1X_2 + 353.757$ | <sup>0.94</sup><sub>1</sub> | 96.456 | <0.0<br>1 |
| 33 | $Y = -0.073X_1^2 + 37.589X_1 - 51.543X_2^2 + 791.949X_2 + 1.078X_1X_2 + 693.764$ | <sup>0.95</sup><sub>6</sub> | 129.27<br>6 | <0.0<br>1 |
| 34 | $Y = -0.072X_1^2 + 36.890X_1 - 51.205X_2^2 + 786.490X_2 + 1.074X_1X_2 + 778.079$ | <sup>0.95</sup><sub>5</sub> | 128.16<br>2 | <0.0<br>1 |
| 35 | $Y = -0.084X_1^2 + 37.846X_1 - 57.559X_2^2 + 1059.350X_2 + 0.859X_1X_2 + 724.847$ | <sup>0.92</sup><sub>8</sub> | 77.111 | <0.0<br>1 |
| 48 | $Y = -0.072X_1^2 + 37.007X_1 - 51.547X_2^2 + 795.226X_2 + 1.073X_1X_2 + 755.152$ | <sup>0.95</sup><sub>5</sub> | 126.91<br>5 | <0.0<br>1 |
| 49 | $Y = -0.074X_1^2 + 38.106X_1 - 51.869X_2^2 + 803.495X_2 + 1.074X_1X_2 + 606.978$ | <sup>0.95</sup><sub>5</sub> | 126.77<br>3 | <0.0<br>1 |
| 50 | $Y = -0.071X_1^2 + 36.684X_1 - 51.349X_2^2 + 791.966X_2 + 1.065X_1X_2 + 783.058$ | <sup>0.95</sup><sub>5</sub> | 128.40<br>7 | <0.0<br>1 |
| 51 | $Y = -0.088X_1^2 + 43.905X_1 - 51.815X_2^2 + 787.471X_2 + 1.154X_1X_2 + 73.936$ | <sup>0.95</sup><sub>5</sub> | 126.07<br>7 | <0.0<br>1 |
| 52 | $Y = -0.069X_1^2 + 36.032X_1 - 51.424X_2^2 + 794.157X_2 + 1.064X_1X_2 + 807.760$ | <sup>0.95</sup><sub>3</sub> | 120.41<br>2 | <0.0<br>1 |
| 53 | $Y = -0.070X_1^2 + 36.173X_1 - 51.847X_2^2 + 800.930X_2 + 1.078X_1X_2 + 812.380$ | <sup>0.95</sup><sub>4</sub> | 123.74<br>3 | <0.0<br>1 |
| 54 | $Y = -0.073X_1^2 + 37.453X_1 - 51.326X_2^2 + 792.352X_2 + 1.069X_1X_2 + 714.406$ | <sup>0.95</sup><sub>3</sub> | 121.06<br>9 | <0.0<br>1 |

## Appendix 2

**Table 2.**
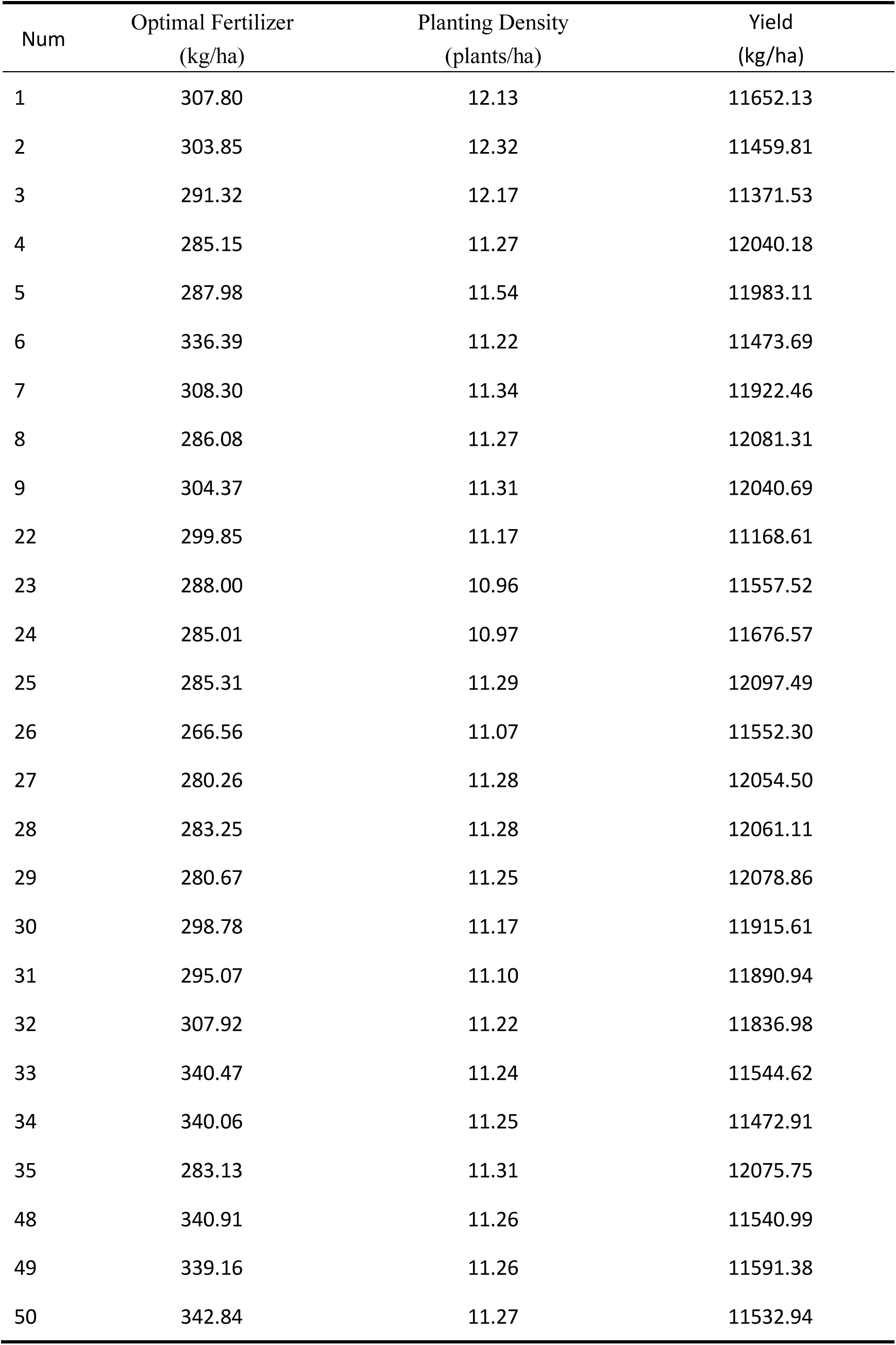
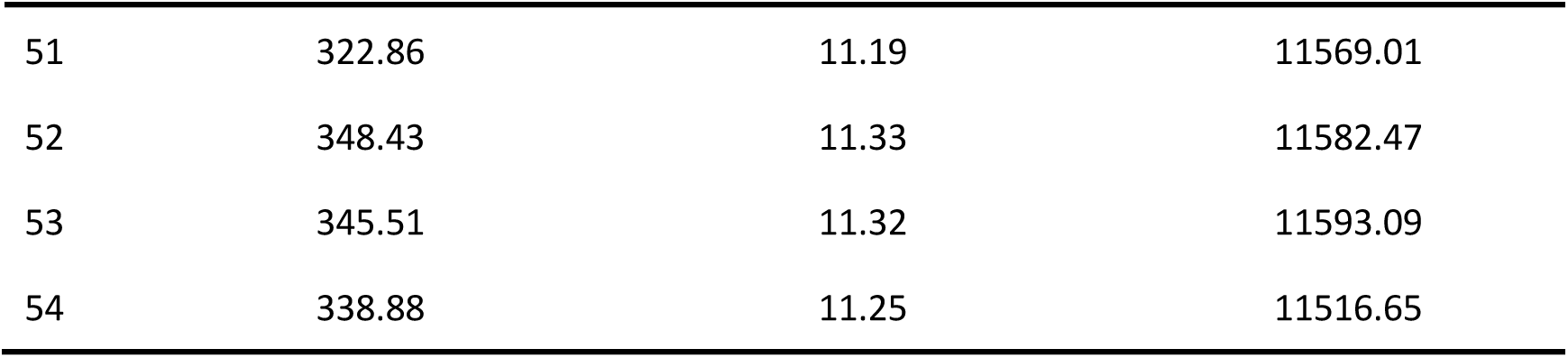
The optimal fertilization amount, planting density, and corresponding yield for planting maize crops on each plot of 290 Farm in 2024.

| Num | Optimal Fertilizer<br>(kg/ha) | Planting Density<br>(plants/ha) | Yield<br>(kg/ha) |
| --- | --- | --- | --- |
| 1 | 307.80 | 12.13 | 11652.13 |
| 2 | 303.85 | 12.32 | 11459.81 |
| 3 | 291.32 | 12.17 | 11371.53 |
| 4 | 285.15 | 11.27 | 12040.18 |
| 5 | 287.98 | 11.54 | 11983.11 |
| 6 | 336.39 | 11.22 | 11473.69 |
| 7 | 308.30 | 11.34 | 11922.46 |
| 8 | 286.08 | 11.27 | 12081.31 |
| 9 | 304.37 | 11.31 | 12040.69 |
| 22 | 299.85 | 11.17 | 11168.61 |
| 23 | 288.00 | 10.96 | 11557.52 |
| 24 | 285.01 | 10.97 | 11676.57 |
| 25 | 285.31 | 11.29 | 12097.49 |
| 26 | 266.56 | 11.07 | 11552.30 |
| 27 | 280.26 | 11.28 | 12054.50 |
| 28 | 283.25 | 11.28 | 12061.11 |
| 29 | 280.67 | 11.25 | 12078.86 |
| 30 | 298.78 | 11.17 | 11915.61 |
| 31 | 295.07 | 11.10 | 11890.94 |
| 32 | 307.92 | 11.22 | 11836.98 |
| 33 | 340.47 | 11.24 | 11544.62 |
| 34 | 340.06 | 11.25 | 11472.91 |
| 35 | 283.13 | 11.31 | 12075.75 |
| 48 | 340.91 | 11.26 | 11540.99 |
| 49 | 339.16 | 11.26 | 11591.38 |
| 50 | 342.84 | 11.27 | 11532.94 |
| 51 | 322.86 | 11.19 | 11569.01 |
| 52 | 348.43 | 11.33 | 11582.47 |
| 53 | 345.51 | 11.32 | 11593.09 |
| 54 | 338.88 | 11.25 | 11516.65 |

## Appendix 3

**S-Table 3.**
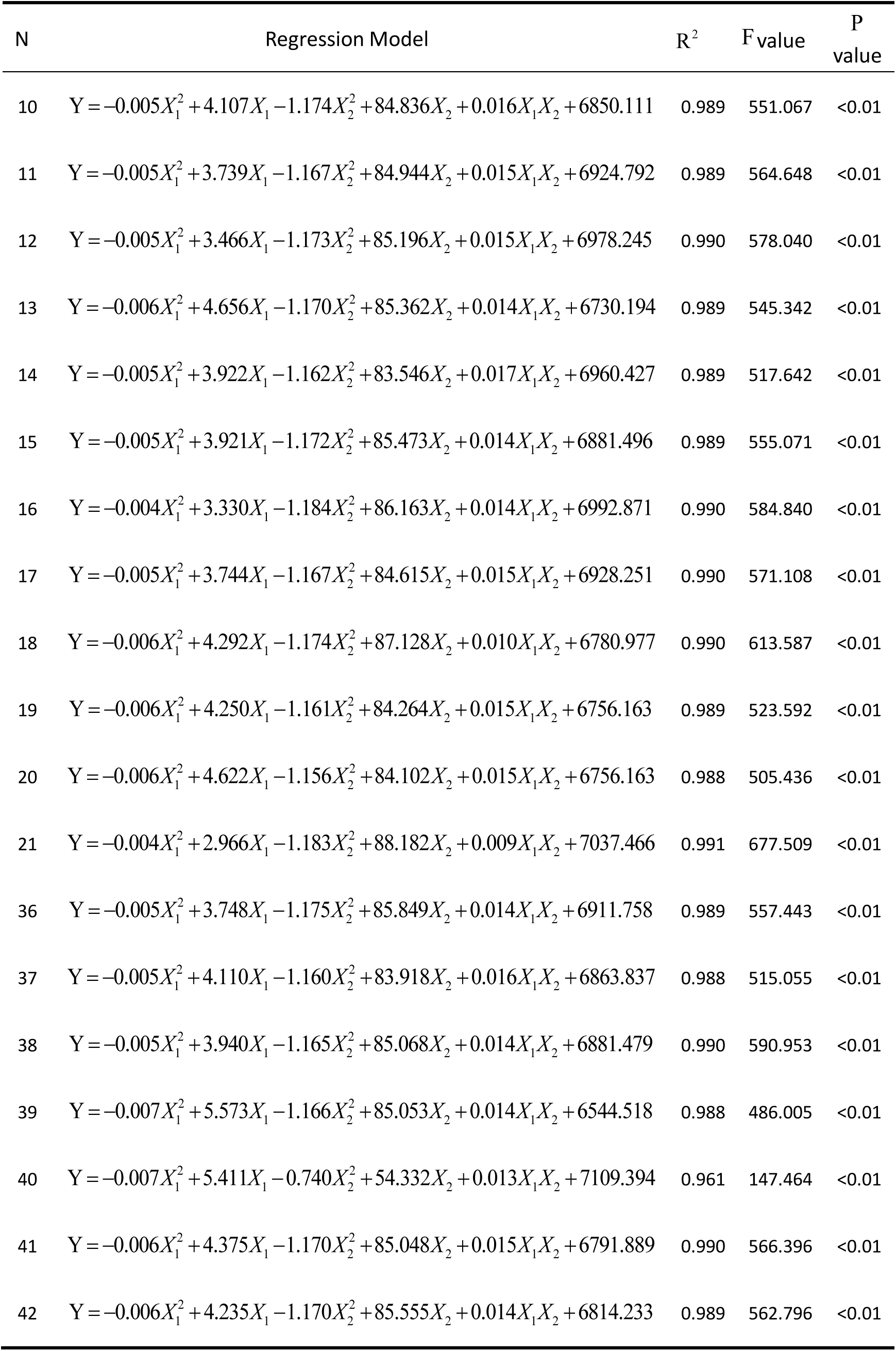
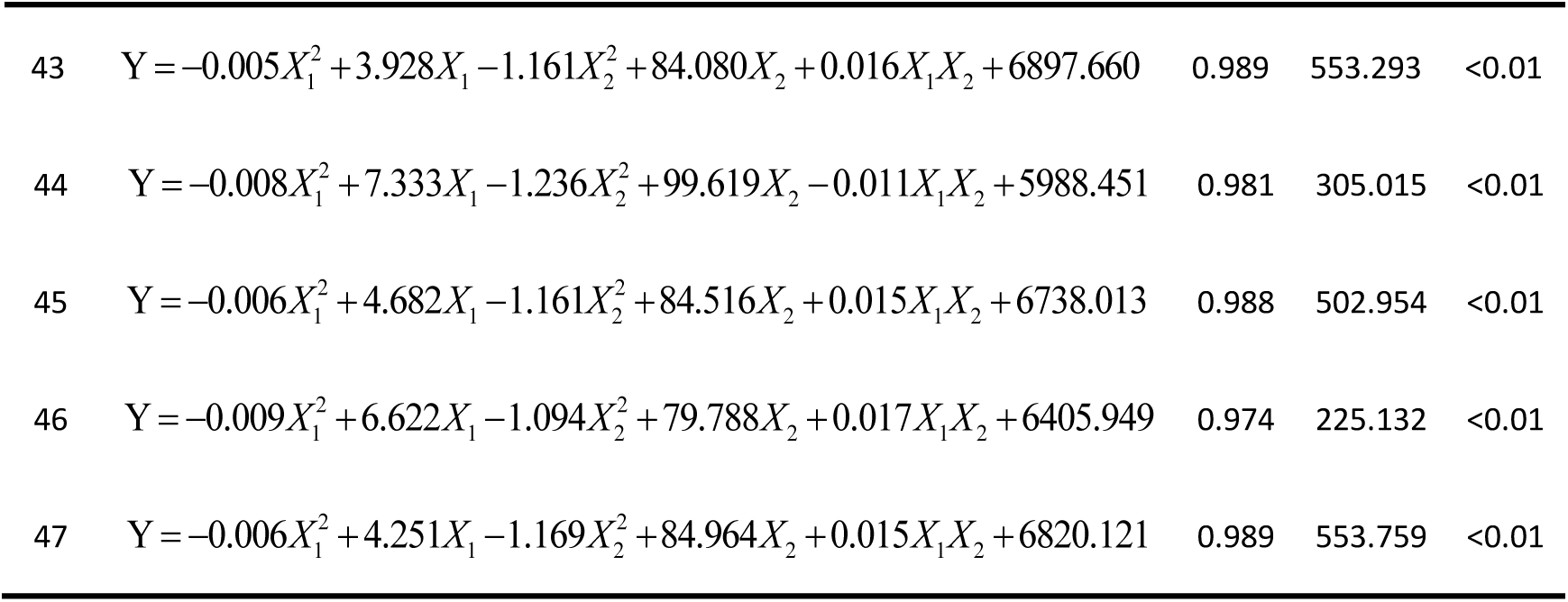
Regression model for unit yield of rice crops planted in 290 Farm in 2024.

Appendix 3 S-Table 3 Regression model for unit yield of rice crops planted in 290 Farm in 2024
| N | Regression Model | R<sup>2</sup> | F value | P value |
| --- | --- | --- | --- | --- |
| 10 | $Y = -0.005X_1^2 + 4.107X_1 - 1.174X_2^2 + 84.836X_2 + 0.016X_1X_2 + 6850.111$ | 0.989 | 551.067 | <0.01 |
| 11 | $Y = -0.005X_1^2 + 3.739X_1 - 1.167X_2^2 + 84.944X_2 + 0.015X_1X_2 + 6924.792$ | 0.989 | 564.648 | <0.01 |
| 12 | $Y = -0.005X_1^2 + 3.466X_1 - 1.173X_2^2 + 85.196X_2 + 0.015X_1X_2 + 6978.245$ | 0.990 | 578.040 | <0.01 |
| 13 | $Y = -0.006X_1^2 + 4.656X_1 - 1.170X_2^2 + 85.362X_2 + 0.014X_1X_2 + 6730.194$ | 0.989 | 545.342 | <0.01 |
| 14 | $Y = -0.005X_1^2 + 3.922X_1 - 1.162X_2^2 + 83.546X_2 + 0.017X_1X_2 + 6960.427$ | 0.989 | 517.642 | <0.01 |
| 15 | $Y = -0.005X_1^2 + 3.921X_1 - 1.172X_2^2 + 85.473X_2 + 0.014X_1X_2 + 6881.496$ | 0.989 | 555.071 | <0.01 |
| 16 | $Y = -0.004X_1^2 + 3.330X_1 - 1.184X_2^2 + 86.163X_2 + 0.014X_1X_2 + 6992.871$ | 0.990 | 584.840 | <0.01 |
| 17 | $Y = -0.005X_1^2 + 3.744X_1 - 1.167X_2^2 + 84.615X_2 + 0.015X_1X_2 + 6928.251$ | 0.990 | 571.108 | <0.01 |
| 18 | $Y = -0.006X_1^2 + 4.292X_1 - 1.174X_2^2 + 87.128X_2 + 0.010X_1X_2 + 6780.977$ | 0.990 | 613.587 | <0.01 |
| 19 | $Y = -0.006X_1^2 + 4.250X_1 - 1.161X_2^2 + 84.264X_2 + 0.015X_1X_2 + 6756.163$ | 0.989 | 523.592 | <0.01 |
| 20 | $Y = -0.006X_1^2 + 4.622X_1 - 1.156X_2^2 + 84.102X_2 + 0.015X_1X_2 + 6756.163$ | 0.988 | 505.436 | <0.01 |
| 21 | $Y = -0.004X_1^2 + 2.966X_1 - 1.183X_2^2 + 88.182X_2 + 0.009X_1X_2 + 7037.466$ | 0.991 | 677.509 | <0.01 |
| 36 | $Y = -0.005X_1^2 + 3.748X_1 - 1.175X_2^2 + 85.849X_2 + 0.014X_1X_2 + 6911.758$ | 0.989 | 557.443 | <0.01 |
| 37 | $Y = -0.005X_1^2 + 4.110X_1 - 1.160X_2^2 + 83.918X_2 + 0.016X_1X_2 + 6863.837$ | 0.988 | 515.055 | <0.01 |
| 38 | $Y = -0.005X_1^2 + 3.940X_1 - 1.165X_2^2 + 85.068X_2 + 0.014X_1X_2 + 6881.479$ | 0.990 | 590.953 | <0.01 |
| 39 | $Y = -0.007X_1^2 + 5.573X_1 - 1.166X_2^2 + 85.053X_2 + 0.014X_1X_2 + 6544.518$ | 0.988 | 486.005 | <0.01 |
| 40 | $Y = -0.007X_1^2 + 5.411X_1 - 0.740X_2^2 + 54.332X_2 + 0.013X_1X_2 + 7109.394$ | 0.961 | 147.464 | <0.01 |
| 41 | $Y = -0.006X_1^2 + 4.375X_1 - 1.170X_2^2 + 85.048X_2 + 0.015X_1X_2 + 6791.889$ | 0.990 | 566.396 | <0.01 |
| 42 | $Y = -0.006X_1^2 + 4.235X_1 - 1.170X_2^2 + 85.555X_2 + 0.014X_1X_2 + 6814.233$ | 0.989 | 562.796 | <0.01 |
| 43 | $Y = -0.005X_1^2 + 3.928X_1 - 1.161X_2^2 + 84.080X_2 + 0.016X_1X_2 + 6897.660$ | 0.989 | 553.293 | <0.01 |
| 44 | $Y = -0.008X_1^2 + 7.333X_1 - 1.236X_2^2 + 99.619X_2 - 0.011X_1X_2 + 5988.451$ | 0.981 | 305.015 | <0.01 |
| 45 | $Y = -0.006X_1^2 + 4.682X_1 - 1.161X_2^2 + 84.516X_2 + 0.015X_1X_2 + 6738.013$ | 0.988 | 502.954 | <0.01 |
| 46 | $Y = -0.009X_1^2 + 6.622X_1 - 1.094X_2^2 + 79.788X_2 + 0.017X_1X_2 + 6405.949$ | 0.974 | 225.132 | <0.01 |
| 47 | $Y = -0.006X_1^2 + 4.251X_1 - 1.169X_2^2 + 84.964X_2 + 0.015X_1X_2 + 6820.121$ | 0.989 | 553.759 | <0.01 |

## Appendix 4

**S-Table 4.**
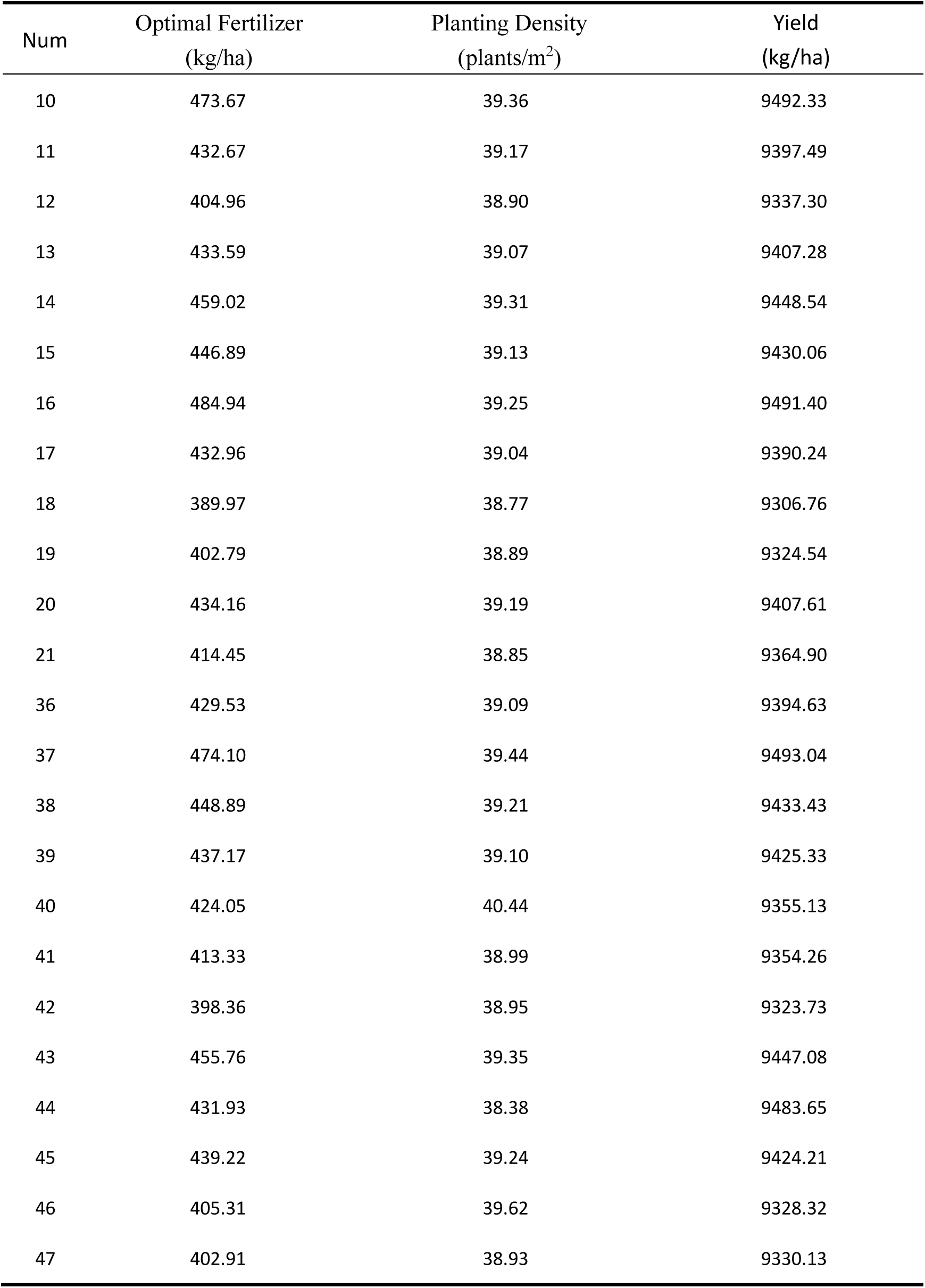
The optimal fertilization amount, planting density, and corresponding yield for planting rice crops on each plot of 290 Farm in 2024.

| Num | Optimal Fertilizer<br>(kg/ha) | Planting Density<br>(plants/m<sup>2</sup>) | Yield<br>(kg/ha) |
| --- | --- | --- | --- |
| 10 | 473.67 | 39.36 | 9492.33 |
| 11 | 432.67 | 39.17 | 9397.49 |
| 12 | 404.96 | 38.90 | 9337.30 |
| 13 | 433.59 | 39.07 | 9407.28 |
| 14 | 459.02 | 39.31 | 9448.54 |
| 15 | 446.89 | 39.13 | 9430.06 |
| 16 | 484.94 | 39.25 | 9491.40 |
| 17 | 432.96 | 39.04 | 9390.24 |
| 18 | 389.97 | 38.77 | 9306.76 |
| 19 | 402.79 | 38.89 | 9324.54 |
| 20 | 434.16 | 39.19 | 9407.61 |
| 21 | 414.45 | 38.85 | 9364.90 |
| 36 | 429.53 | 39.09 | 9394.63 |
| 37 | 474.10 | 39.44 | 9493.04 |
| 38 | 448.89 | 39.21 | 9433.43 |
| 39 | 437.17 | 39.10 | 9425.33 |
| 40 | 424.05 | 40.44 | 9355.13 |
| 41 | 413.33 | 38.99 | 9354.26 |
| 42 | 398.36 | 38.95 | 9323.73 |
| 43 | 455.76 | 39.35 | 9447.08 |
| 44 | 431.93 | 38.38 | 9483.65 |
| 45 | 439.22 | 39.24 | 9424.21 |
| 46 | 405.31 | 39.62 | 9328.32 |
| 47 | 402.91 | 38.93 | 9330.13 |

## Appendix 5

**S-Table 5.** Regression model for unit yield of soybean crops planted in 290 Farm in 2024.

Appendix 5 S-Table 5 Regression model for unit yield of soybean crops planted in 290 Farm in 2024
| N | Regression Model | R<sup>2</sup> | F value | P value |
| --- | --- | --- | --- | --- |
| 1 | $Y = -0.006X_1^2 + 1.912X_1 - 0.248X_2^2 + 15.842X_2 + 2432.803$ | 0.923 | 92.499 | <0.01 |
| 2 | $Y = -0.006X_1^2 + 1.793X_1 - 0.244X_2^2 + 15.591X_2 + 2450.092$ | 0.940 | 121.59<br>1 | <0.01 |
| 3 | $Y = -0.006X_1^2 + 1.847X_1 - 0.256X_2^2 + 16.173X_2 + 2435.484$ | 0.918 | 56.842 | <0.01 |
| 4 | $Y = -0.007X_1^2 + 2.235X_1 - 0.255X_2^2 + 16.277X_2 + 2498.357$ | 0.926 | 96.961 | <0.01 |
| 5 | $Y = -0.006X_1^2 + 1.840X_1 - 0.240X_2^2 + 15.297X_2 + 2442.832$ | 0.931 | 104.90<br>7 | <0.01 |
| 6 | $Y = -0.007X_1^2 + 2.444X_1 - 0.288X_2^2 + 19.436X_2 + 2239.309$ | 0.959 | 183.09<br>8 | <0.01 |
| 7 | $Y = -0.007X_1^2 + 2.222X_1 - 0.227X_2^2 + 14.803X_2 + 2416.563$ | 0.919 | 88.148 | <0.01 |
| 8 | $Y = -0.006X_1^2 + 2.171X_1 - 0.225X_2^2 + 14.728X_2 + 2419.726$ | 0.922 | 91.243 | <0.01 |
| 9 | $Y = -0.006X_1^2 + 2.100X_1 - 0.221X_2^2 + 14.611X_2 + 2422.811$ | 0.924 | 94.894 | <0.01 |
| 22 | $Y = -0.006X_1^2 + 2.711X_1 - 0.305X_2^2 + 24.070X_2 - 0.012X_1X_2 + 2115.156$ | 0.982 | 319.78<br>0 | <0.01 |
| 23 | $Y = -0.004X_1^2 + 1.165X_1 - 0.663X_2^2 + 44.540X_2 + 0.034X_1X_2 + 1309.337$ | 0.975 | 234.92<br>5 | <0.01 |
| 24 | $Y = -0.004X_1^2 + 1.130X_1 - 0.811X_2^2 + 50.678X_2 + 0.030X_1X_2 + 1348.392$ | 0.977 | 249.66<br>4 | <0.01 |
| 25 | $Y = -0.006X_1^2 + 2.133X_1 - 0.251X_2^2 + 15.915X_2 + 2413.974$ | 0.932 | 106.07<br>7 | <0.01 |
| 26 | $Y = -0.007X_1^2 + 2.332X_1 - 0.278X_2^2 + 17.719X_2 + 2366.529$ | 0.924 | 94.426 | <0.01 |
| 27 | $Y = -0.007X_1^2 + 2.273X_1 - 0.248X_2^2 + 15.769X_2 + 2404.675$ | 0.916 | 85.004 | <0.01 |
| 28 | $Y = -0.007X_1^2 + 2.220X_1 - 0.236X_2^2 + 15.184X_2 + 2416.148$ | 0.909 | 77.106 | <0.01 |
| 29 | $Y = -0.007X_1^2 + 2.188X_1 - 0.251X_2^2 + 15.762X_2 + 2419.540$ | 0.915 | 83.010 | <0.01 |
| 30 | $Y = -0.006X_1^2 + 2.201X_1 - 0.239X_2^2 + 15.443X_2 + 2412.757$ | 0.930 | 102.41<br>1 | <0.01 |
| 31 | $Y = -0.006X_1^2 + 2.219X_1 - 0.224X_2^2 + 14.777X_2 + 2409.855$ | 0.924 | 94.578 | <0.01 |
| 32 | $Y = -0.006X_1^2 + 2.168X_1 - 0.243X_2^2 + 15.709X_2 + 2399.212$ | 0.937 | 116.23<br>7 | <0.01 |
| 33 | $Y = -0.007X_1^2 + 2.451X_1 - 0.293X_2^2 + 19.717X_2 + 2230.837$ | 0.960 | 187.40<br>3 | <0.01 |
| 34 | $Y = -0.007X_1^2 + 2.416X_1 - 0.291X_2^2 + 19.661X_2 + 2233.171$ | 0.959 | 181.16<br>5 | <0.01 |
| 35 | $Y = -0.007X_1^2 + 2.164X_1 - 0.249X_2^2 + 15.632X_2 + 2419.579$ | 0.912 | 80.499 | <0.01 |
| 48 | $Y = -0.006X_1^2 + 2.318X_1 - 0.312X_2^2 + 20.696X_2 + 2224.537$ | 0.965 | 216.79<br>9 | <0.01 |
| 49 | $Y = -0.007X_1^2 + 2.355X_1 - 0.282X_2^2 + 19.275X_2 + 2241.943$ | 0.962 | 198.23<br>1 | <0.01 |
| 50 | $Y = -0.006X_1^2 + 2.318X_1 - 0.313X_2^2 + 20.828X_2 + 2224.314$ | 0.968 | 235.09<br>9 | <0.01 |
| 51 | $Y = -0.006X_1^2 + 2.412X_1 - 0.294X_2^2 + 18.962X_2 + 2250.361$ | 0.946 | 135.36<br>1 | <0.01 |
| 52 | $Y = -0.007X_1^2 + 2.404X_1 - 0.279X_2^2 + 19.070X_2 + 2233.841$ | 0.957 | 172.46<br>5 | <0.01 |
| 53 | $Y = -0.006X_1^2 + 2.285X_1 - 0.287X_2^2 + 19.469X_2 + 2244.940$ | 0.960 | 186.90<br>7 | <0.01 |
| 54 | $Y = -0.007X_1^2 + 2.418X_1 - 0.279X_2^2 + 19.019X_2 + 2247.460$ | 0.958 | 175.97<br>9 | <0.01 |

## Appendix 6

**S-Table 6.** The optimal fertilization amount, planting density, and corresponding unit yield for planting soybean crops on each plot of 290 Farm in 2024.

| Num | Optimal Fertilizer<br>(kg/ha) | Planting Density<br>(plants/m<sup>2</sup>) | Yield<br>(kg/ha) |
| --- | --- | --- | --- |
| 1 | 159.33 | 31.94 | 2838.19 |
| 2 | 149.42 | 31.95 | 2833.10 |
| 3 | 153.92 | 31.59 | 2833.06 |
| 4 | 159.64 | 31.92 | 2936.50 |
| 5 | 153.33 | 31.87 | 2827.65 |
| 6 | 174.57 | 33.74 | 2780.55 |
| 7 | 158.71 | 32.61 | 2834.23 |
| 8 | 180.92 | 32.73 | 2857.13 |
| 9 | 175.00 | 33.06 | 2848.06 |
| 22 | 190.20 | 35.72 | 2802.83 |
| 23 | 323.65 | 41.89 | 2430.72 |
| 24 | 277.67 | 36.38 | 2427.11 |
| 25 | 177.75 | 31.70 | 2855.82 |
| 26 | 166.57 | 31.87 | 2843.09 |
| 27 | 162.36 | 31.79 | 2839.86 |
| 28 | 158.57 | 32.17 | 2836.39 |
| 29 | 156.29 | 31.40 | 2837.97 |
| 30 | 183.42 | 32.31 | 2864.07 |
| 31 | 184.92 | 32.98 | 2858.73 |
| 32 | 180.67 | 32.32 | 2848.94 |
| 33 | 175.07 | 33.65 | 2777.09 |
| 34 | 172.57 | 33.78 | 2773.73 |
| 35 | 154.57 | 31.39 | 2832.17 |
| 48 | 193.17 | 33.17 | 2791.63 |
| 49 | 168.21 | 34.18 | 2769.38 |
| 50 | 193.17 | 33.27 | 2794.68 |
| 51 | 201.00 | 32.25 | 2798.51 |
| 52 | 171.71 | 31.18 | 2766.11 |
| 53 | 190.42 | 33.92 | 2792.67 |
| 54 | 172.71 | 34.08 | 2780.40 |

